# LanTERN: a fluorescent sensor that specifically binds lanthanides

**DOI:** 10.1101/2023.09.26.559553

**Authors:** Ethan Jones, Yang Su, Chris Sander, Quincey A. Justman, Michael Springer, Pamela A. Silver

## Abstract

Lanthanides, a series of fifteen f-block elements, are crucial in modern technology, and their purification by conventional chemical means comes at a significant environmental cost. Synthetic biology offers promising solutions. However, the lack of biochemical tools to measure lanthanide binding is a bottleneck to progress. Here, we introduce LanTERN, a lanthanide-responsive fluorescent protein rationally engineered from the lanmodulin-binding protein, LanM. LanTERN was designed based on GCaMP, a genetically encoded calcium indicator that couples the ion binding of four EF hand motifs to increased GFP fluorescence. We engineered seven mutations across the parent construct’s four EF hand motifs to switch specificity from calcium to lanthanides. The resulting protein, LanTERN, directly converts the binding of 10 measured lanthanides to 14-fold or greater increased fluorescence. LanTERN development opens new avenues for creating improved lanthanide-binding proteins and bio-sensing systems.

## Introduction

Modern technology relies on lanthanides, a series of fifteen f-block elements. Lanthanides have similar physicochemical properties and co-occur within the Earth’s crust, but must be separated for use in technological applications. Separation of lanthanides is environmentally costly and typically requires tens or hundreds of stages of solvent extraction.^1^ Given these elements’ critical importance, new, cost-effective, environmentally friendly methods of lanthanide separation are needed.

Natural microbes contain protein-based machinery that bind individual lanthanides with varying affinities. For example, some methanotrophic organisms preferentially import light lanthanides and use them as co-factors for alcohol dehydrogenases.^2^ Many of these organisms also contain lanmodulins (LanMs), a family of proteins with a structure similar to calmodulin. These LanMs use distinct variants of the EF hand motif to specifically bind to lanthanides, with a slight preference for the lighter lanthanides.^2–4^ Recently, LanMs were used to help separate mixtures of lanthanides such as neodymium and dysprosium.^3,4^ Synthetic biology offers the potential to develop enhanced molecules and organisms that bind to, discriminate between, and thereby separate different lanthanide elements.

Synthetic biology is able to engineer proteins with have novel or improved functionality such as binding, provided that those functions can be measured. However, our ability to measure the binding of lanthanides to proteins is limited. While one FRET-based sensor detects lanthanides by conformational change of the native LanM protein, there is currently no published single fluorophore sensor that directly transduces lanthanide ion binding into a fluorescent output.^5^ Here, we describe a lanthanide sensor, LanTERN, that directly couples lanthanide binding to a change in green fluorescent protein (GFP) signal.

## Results

We designed our fluorescent sensor based on GCaMP, a genetically encoded calcium indicator. GCaMP has been extensively optimized as a reporter of intracellular calcium concentration in the context of live-cell imaging.^6^ We wanted to switch the specificity of GCaMP from calcium to lanthanides to create a fluorescent sensor that could be used to measure the binding of lanthanides in vitro.

GCaMP is a fusion protein that comprises an N-terminal calmodulin-binding peptide (CBP), a circularly permuted green fluorescent protein (cpGFP), and a C-terminal calmodulin. Calmodulin, in turn, is comprised of four 12 amino acid calcium-binding motifs (EF hands) separated by 24- or 25-amino acid linkers.^7^ In the presence of calcium, calmodulin undergoes a marked conformational change, exposing an internal binding site for CBP to bind in *cis*. This intramolecular binding increases the brightness of GFP by excluding water from its chromophore.^8^

The conformational shift by calmodulin is driven by the binding of calcium ions to its four EF hand motifs. To switch GCaMP’s specificity from calcium to lanthanides, we used the EF hands of *Methylorubrum extorquens* LanM (*mex*-LanM) to construct a lanthanide responsive super folder GCaMP variant by combining an M13 CBP and a circularly permutated super folder GFP with a chimeric *Rattus norvegicus* calmodulin in which each of the 4 EF hands was replaced with the corresponding EF hand from *mex*-LanM **(Figure 1a, middle, and Figure 1b, blue)**.

**Figure 1.**
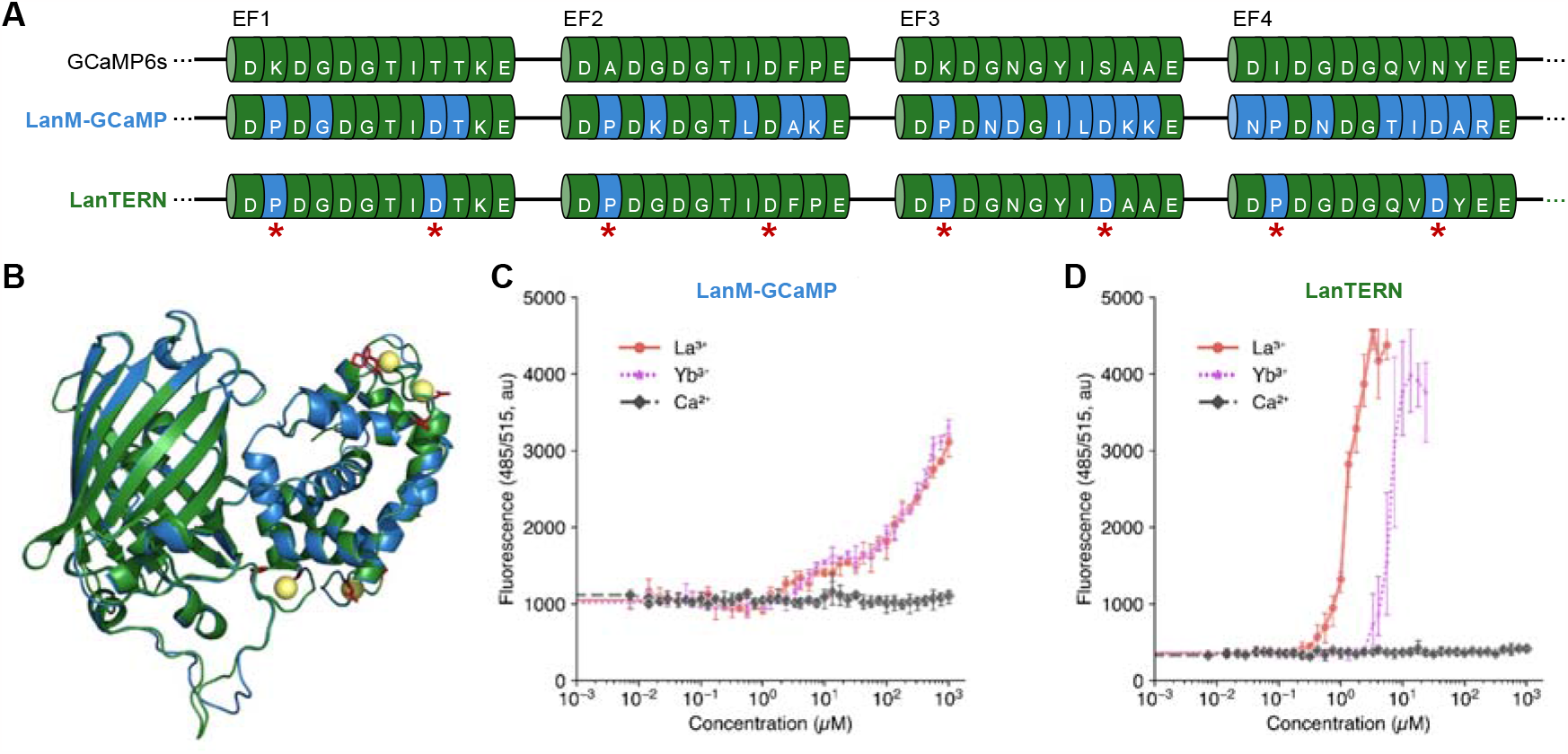
Rational Engineering of EF Hand motifs convert GCaMP into a lanthanide sensor: **a.** Sequences of EF hands 1, 2, 3, and 4 of GCaMP, LanM-GCaMP, and LanTERN. Amino acids identical to GCaMP are green; amino acids identical to LanM-GCaMP are blue; intervening linkers (not to scale) are depicted as lines. Red stars indicate amino acid sidechains shown as red sticks in panel B **b.** Overlaid models of metal-bound LanM-GCaMP (blue) and LanTERN (green). Prolines at EF hand position 2 and putative lanthanide-binding aspartates at EF hand position 9 are shown as red sticks. **c.** Fluorescence measurements of 500nM LanM-GCaMP (see Fig. 1A, middle and Fig. 1B, blue) in the presence of varying calcium, lanthanum, and ytterbium concentrations. Points and error bars represent the mean and standard deviation of three technical replicates from the same protein purification and working dilution. Graphs of two additional protein purifications can be found in **Supplemental Figure 3**. **d.** Fluorescence measurements of 500nM LanTERN (see Fig. 1A, bottom and Fig. 1B, green) in varying lanthanum, ytterbium, and calcium concentrations. Points and error bars represent the mean and standard deviation of three technical replicates from the same protein purification and working dilution. Graphs of two additional protein purifications can be found in **Supplemental Figure 4**.

We expressed this construct, termed LanM-GCaMP, in *Escherichia coli* (*E. coli*), purified it using Ni-NTA chromatography (see **Supplemental Methods**), and measured its dose response *in vitro* to lanthanum, the lightest lanthanide, ytterbium, the heaviest non d-block lanthanide, and calcium. Installation of the EF hands from LanM increased brightness of the GFP in the presence of lanthanides (approximately 10μM-1mM) but not in the presence of calcium **(Figure 1c)**. This stands in contrast to the K_d_ of *mex-*LanM, which is reported to be in the picomolar range.^3^ We reasoned that the relatively weak response might have been due to the orientation of the hands in the calmodulin backbone; EF hands are paired in the ion-bound conformation, and LanM and calmodulin differ in their arrangement.^3,7^ We tested three alternative hand arrangements to mimic the native arrangement in *mex*-LanM but did not observe improved performance. **(Supplemental Figure 1)**.

We reasoned that we could improve the performance of the sensor by installing the features of the LanM EF hands that are thought to confer lanthanide selectivity^3,4^ in the native calmodulin EF hands. In *mex*-LanM, the proline residue in the second position of each EF hand is critical to specificity. Mutating these proline residues ablates *mex*-LanM’s specificity for lanthanides vs. calcium^3^; they are conserved across LanMs.^4^ Therefore, we mutated the second position of each EF hand to a proline. We also mutated each of the first four metal contacting residues (EF hand residues 1,3,5, and 9) to aspartic acid because these residues directly coordinate lanthanides in *mex*-LanM.^3,4,93,4,9^ The resulting construct was termed LanTERN, for <u>lan</u>thanide-<u>t</u>uned <u>E</u>F hand <u>r</u>eporter fluoresce<u>n</u>t (**Figure 1a, bottom, Fig 1b, green**).

We measured the dose-response of LanTERN to lanthanides. The concentration of lanthanides that yielded the maximal response (Ln_max_) varied among lanthanides (0.5μM-10μM). Above Ln_max_ we observed non monotonic behavior in the fluorescence (**Supplemental Figure 6**). Therefore we defined LanTERN’s dynamic range as [Ln]<[Ln_max_] and restricted our characterization and analysis to this range.

Within LanTERN’s dynamic range, we calculated EC_50_s for LanTERN to be 976nM for lanthanum and 4.71μM for ytterbium, with no measurable response to calcium **(Supplemental Table 1)**. This is an improvement of greater than two orders of magnitude over the LanM-GCaMP. LanTERN’s lanthanide-dependent increase in fluorescence vs baseline also increased over LanM-GCaMP’s (e.g. >10-fold change vs. ∼3.5-fold change in response to lanthanum) **(Figure 1d)**. To confirm the importance of the proline residue in the second position of each engineered EF hand, we created a LanTERN variant where the second position proline was back-mutated to the cognate amino acid found in wild type calmodulin. As expected, these mutations reduced the sensor’s response to lanthanides by approximately 10-fold and restored its response to calcium **(Supplemental Figure 2)**.

Finally, we characterized LanTERN’s response to 10 lanthanides that span the atomic weight of this class. LanTERN responds to all lanthanides tested: we observed a 14-fold or greater lanthanide dependent increase in fluorescence vs. baseline **(Figure 2)**. The sensor exhibited binding preferences similar to *mex*-LanM, generally responding at lower concentrations of the lighter lanthanides **(Table 1)**.^3,4^ The difference in EC_50_ values for the lightest and heaviest lanthanides tested differed by approximately 4-fold.

**Figure 2:**
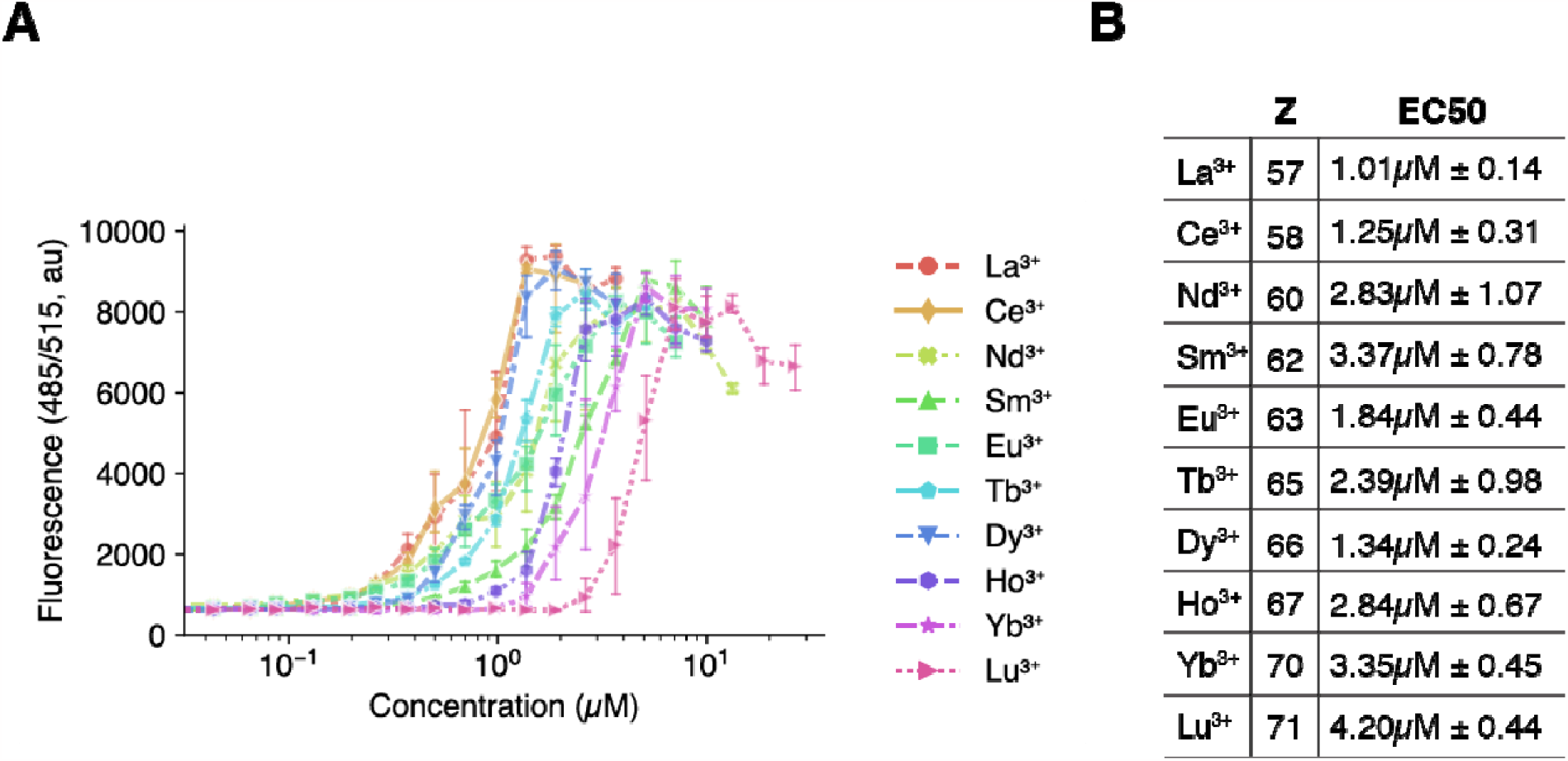
LanTERN responds to all lanthanides. **a.** Fluorescence measurements of 500nM LanTERN in the presence of varying concentrations of lanthanides listed in order of atomic mass. Points and error bars represent the mean and standard deviation of three technical replicates from the same protein purification and working dilution. Lines represent a linear interpolation between points. Graphs of two additional protein purifications can be found in **Supplemental Figure 8**. **b.** Table of calculated EC_50_s of LanTERN in response to lanthanides. Lanthanides are shown in order of atomic number (Z). Values represent the mean of three independent protein purifications of LanTERN, ± represents the standard deviation of this mean. Values for the individual protein purifications can be found in **Supplemental Table 1**.

## Discussion

In this study, we report the construction of LanTERN, a lanthanide-responsive fluorescent protein. We rationally engineered EF hand motifs to build a fluorescent protein sensor with switched specificity for lanthanides vs. calcium. This capability opens new avenues for the creation of improved lanthanide-binding proteins. LanTERN could be used as a sensing tool in directed evolution studies to identify mutations in EF hands that increase selectivity and affinity for specific lanthanides. Lanthanide binders with higher specificity could be used in synthetic biological approaches for separating lanthanides. Alternatively, LanTERN could be used to develop engineered calmodulin domains for bio-sensing systems such as lanthanide-responsive transcriptional regulators. Such systems might enable the creation of organisms that respond to the presence of lanthanides and assist in the extraction and separation of lanthanides.

## Methods

Detailed protocols for all methods used in this report are given in the supplemental information.

## Supporting Information

### SI1.pdf

- Supplemental methods and protocols for experiments and analysis performed in this report
- Supplemental Figures:
  ∘ Figure S1: La, Yb, and Ca dose-response of LanM-GCaMP variants with different LanM hand ordering.
  ∘ Figure S2: La, Yb, and Ca dose-response of LanTERN with back mutations in the second position of each EF hand.
  ∘ Figure S3: La, Yb, and Ca dose-response of LanM-GCaMP including all 3 independent protein purifications.
  ∘ Figure S4: La, Yb, and Ca dose-response of LanTERN including all 3 independent protein purifications.
  ∘ Figure S5: La, Yb, and Ca dose-response of LanM-GCaMP variants with different LanM hand orderings including all 3 independent protein purifications.
  ∘ Figure S6: La, Yb, and Ca dose-response of LanTERN showing non-monotonic dynamics outside of Ln_max_
  ∘ Figure S7: La, Yb, and Ca dose-response of LanTERN with back mutations in the second position of each EF hand including all 3 independent protein purifications.
  ∘ Figure S8: Dose-response of LanM-GCaMP to 10 lanthindes including all 3 independent protein purifications.
- Supplemental Tables
  ∘ Table ST1: Calculated EC50 values for LanTERN from figure 1D
  ∘ Table ST2: Oligonucleotides used in this study
  ∘ Table ST3: Constructs used in this study
- Supplemental Materials: List of catalog numbers for materials used in this report S2.zip

### S2.zip

- Raw and processed data from each experiment in this report, jupyter notebooks used to analyze and plot this data, annotated genbank files for constructs used in this report, .pdb files of structures shown in Figure 2B

## Author Information

### Author Contribution

EMJ, PAS, QAJ, and MS conceptualized project direction; EMJ conceptualized, designed, and cloned sensors, planned and conducted experiments, and analyzed data. YS constructed the protein structure models of LanM-GCaMP and LanTERN. EMJ, QAJ, and YS created figures; EMJ, PAS, and QAJ wrote and edited manuscript. All authors approved the manuscript.

### Notes

The authors declare no competing financial interest.

## Acknowledgement

pRSET sfGCaMP6s-T78H was a gift from Wolf Frommer (Addgene plasmid #100023). The authors thank the Laboratory of Systems Pharmacology at Harvard Medical School for access to their equipment and Neil Dalvie for his helpful comments on the manuscript.

This material is based upon work supported by the National Science Foundation Graduate Research Fellowship under Grant No. DGE 2140743. This work was supported by funds from the MITRE Corporation, the Wyss Institute for Biologically Inspired Engineering and the Synthetic Biology HIVE at Harvard Medical School

