## Supplemental Information 1 for "LanTERN: a fluorescent sensor that specifically binds lanthanides"

### Supplemental Methods

#### Design of Constructs and Cloning

The constructs used in this study (see Supplemental Table 3) were all constructed using double-stranded gene fragments (Twist and IDT). An agar stab of pRSET sfGCaMP6s-T78H was obtained from addgene (Plasmid #100023). The stab was restruck, inoculated in selective LB, miniprepped, and the plasmid sequenced using whole plasmid long-read sequencing (Primordium). Each plasmid in Supplemental Table 3 was constructed in a two-fragment isothermal assembly using New England Biolabs' (NEB) NEBuilder® HiFi DNA Assembly Master Mix.

A PCR amplicon of the pRSET sfGCaMP6s-T78H vector generated using Q5® High-Fidelity 2X Master Mix (NEB) with the primers EMJ_P088 and EMJ_P089 (see Supplemental Table 2)[i] was used as the vector fragment in each assembly. The presence of a single band was confirmed via agarose gel electrophoresis, the PCR treated with DpnI[ii] (NEB) to remove background, and then purified with a PCR purification column (Zymo). DNA concentration was then determined using a NanoDrop One (ThermoFisher).

Then, each isothermal assembly reaction was performed according to a miniaturized version of the manufacturer's protocol. Each reaction had a total volume of 5µL and used 0.03 picomoles of vector and 0.09 picomoles of the corresponding gene fragment (see Supplemental Table 2). The reactions were incubated at 50°C for 1 hour, and then 0.5µL of each reaction was transformed into 10µL of high efficiency chemically competent 5-alpha *E. coli* (NEB) following the manufacturer's protocol. 50µL was plated on ampicillin or carbenicillin-containing plates (100µg/mL), individual colonies were picked, miniprepped (Epoch Life Sciences)[iii], and sequence confirmed using whole plasmid long-read sequencing (Primordium).

#### Protein Expression and Purification

Miniprepped constructs were transformed into chemically competent NiCo21(DE3) *E. coli* from New England Biolabs using the manufacturers protocol and plated on carbenicillin or ampicillin-containing plates (100µg/mL). For the results presented in the paper, 3 distinct colonies (biological replicates) of each construct were independently picked into 200mL of ZYM-5052 autoinduction media[1] containing 100µg/mL of carbenicillin. Cultures were grown in 1000mL baffled flasks in a shaking incubator set to 250RPM at 30°C[iv] for 36-48 hours[v].

---

[i] 98°C denaturation for 30 seconds, 27 cycles of 98°C 10 seconds, 64°C 30 seconds, and 72°C for 2 minutes, then a final extension at 72°C for 2 minutes

[ii] Add 1x CutSmart®, incubate at 37°C for 1 hour and 80°C heat kill for 20 minutes

[iii] Columns from Epoch Life Science were used, but buffers and protocol were those of standard Qiagen miniprep. Some buffers were prepared from scratch rather than purchased.

[iv] LanTERN variants have all proven stable at 37°C, although as is typical protein yield was higher at lower temperatures for longer periods, likely due to increased oxygen solubility

[v] 2 overnights

Cells were centrifuged at ~4000 RCF and resuspended in ~10-15mL buffer chemically identical[vi] to Takara His60 Ni Superflow equilibration buffer (50 mM sodium phosphate, 300 mM sodium chloride, 20 mM imidazole; pH 7.4). Cells were transferred to 50mL conical tubes (Corning) and lysed by probe sonication on ice for approximately 3-4 minutes. The lysate was clarified by centrifugation at 4337 RCF. Complete lysis of the pellet was checked by inspecting the pellet briefly under a blue light box and ensuring that the pellet had minimal fluorescence compared to the medium.

Clarified lysate (supernatant) was transferred to a 10mL disposable column (ThermoFisher) containing 1mL of chitin resin (NEB) that had been washed with the above equilibration buffer (3x20mL). Columns were then incubated with end-over-end rotation at 4°C for 30 minutes. Columns were then drained, and the flow through retained. The chitin resin was then washed with ~10ml of equilibration buffer, which was combined with the flow through.The combined wash and flow-through were added to new columns containing 2mL of equilibrated[vii] Takara His60 Ni Superflow resin[viii]. Columns were incubated with end-over-end rotation at 4°C overnight. The next day, columns were drained, washed once with 10mL of equilibration buffer, washed once with 10-15mL of Takara His60 Ni Superflow wash buffer (50 mM sodium phosphate, 300 mM sodium chloride, 40 mM imidazole; pH 7.4) and then eluted using 15mL[ix] Takara His60 Ni Superflow elution buffer (50 mM sodium phosphate, 300 mM sodium chloride, 300 mM imidazole; pH 7.4)[x]. Eluted proteins were transferred to a 10,000kDa MW cut-off protein concentrator (Millipore) and concentrated to a 3mL final volume. Proteins were then exchanged into Buffer C (30 mM MOPS, 100 mM KCl, pH 7.2) from Mattocks et al. (2019)[2] using an Econo-Pac 10DG Desalting Column (Bio-Rad). Then, protein concentration was measured by performing the microwell BCA protocol from the Pierce™ BCA Protein Assay Kit (ThermoFisher) according to the manufacturer's protocol.[xi] Three 20μL samples were measured for each protein, and three technical replicates of each member of the BSA standard curve (in buffer C)[xii] were used on each plate. Measurements between technical replicates were in close agreement, and the mean of the three replicates was used alongside a fit to the standard curve to determine the concentration. Based on these concentrations, 80% glycerol and buffer C were added to the protein to achieve a final concentration of either 5μM or 10μM protein with a 20% glycerol concentration. Raw data of the BCAs, standard curves, and dilution calculations can be found in the experimental information supplement. Proteins were then aliquoted into single-use tubes and snap-frozen in liquid nitrogen before being transferred to storage in a -80°C freezer.

---

[vi] i.e. Prepared from scratch rather than purchased

[vii] Resin is equilibrated off column by spinning down for 2 minutes at 100RCF and then resuspending in equilibration buffer 2-3 times.

[viii] Resin was always fresh and not reused

[ix] 15mL elution was done in two stages, 10mL and then an additional 5mL

[x] Both buffers again prepared from scratch rather than purchased

[xi] https://web.archive.org/web/20230926153642/https://assets.thermofisher.com/TFS-Assets%2FLSG%2Fmanuals%2FMAN0011430_Pierce_BCA_Protein_Asy_UG.pdf

[xii] See ThermoFisher protocol

#### In vitro experiments

For in vitro experiments, protein aliquots were first thawed in a heatblock set to 37°C and then placed onto ice. Then, working solutions were made in Buffer C to a final concentration of 500nM protein and 2% glycerol. Each 384-well plate (Greiner Bio-One) well was filled with 50µL of the working solution, covered, and spun down briefly. 99mM stocks of lanthanides (Alfa Aesar, ThermoFisher, Sigma Aldrich) in 0.1% Triton-X and distilled water were prepared by mixing 990µL of 100mM lanthanide nitrate stocks in distilled water with 10µL of 10% Triton X-100. Lanthanide solutions were then dispensed using a D300e (HP) dispenser, with each plate containing three technical replicates for each concentration/condition. After dispensing, plates were covered, briefly spun down, and measured at ambient temperature on a Bio-Tek H1 plate reader with an emission wavelength of 480nm and excitation wavelength of 515nm. Data from each experiment was processed using Python. In brief, data from each plate reader experiment was combined with its associated concentration data, and dose-response curves were generated by taking all data up until saturation, which was defined as the point at which the mean of a given experiment was no longer increasing for at least three consecutive data points, given that there had been an at least 1.5x fold change. That data was then saved and used to generate plots. EC50s were calculated by calculating the maximal fluorescence value and associated concentration, finding the nearest two points to the half-maximal fluorescence value, and performing a linear interpolation to determine the concentration value associated with the value (see calculate_half_max.ipynb in supplemental information 2). The complete raw data from the instruments (including dispense information from the D300e), as well as intermediates used for the construction of plots, can be found in the supplementary files. Python notebooks containing the code used to generate the plots can also be found in supplementary files. The Python environment used for these notebooks was the first author's personal Python conda environment; a yaml export of the conda environment's installed packages is provided as plotting_enviroment.yml.[xiii]

#### Structural Modeling

Structural models of LanTERN and LanM-GCaMP hands were built with Modeller. PDB 3ek7, 3ek8, 3ekh, 7aug, 6zsm, 6zsn, 6tv7, 3o77, 3o78, 3evr, 3evu, 3evv, 6ya9, 3sg2, 3sg3, 3sg4, 3sg5, 3sg6, 3sg7, 4ik1, 4ik3, 4ik4, 4ik5, 4ik8, 4ik9, 3u0k, 6xu4, 3wlc, 3wld, and 4i2y were used as templates for modeling LanTERN and GCaMP with LanM hands. The EF-hand loops of PDB 8fns and 6mi5 were also used as templates for modeling GCaMP with LanM hands.[3]

---

[xiii] This was performed on an M1 Mac (Ventura 13.1). Multiple unneeded packages are present as the environment was used for other unrelated analyses. The authors apologize for the additional and unnecessary dependencies.

### Supplemental Figures

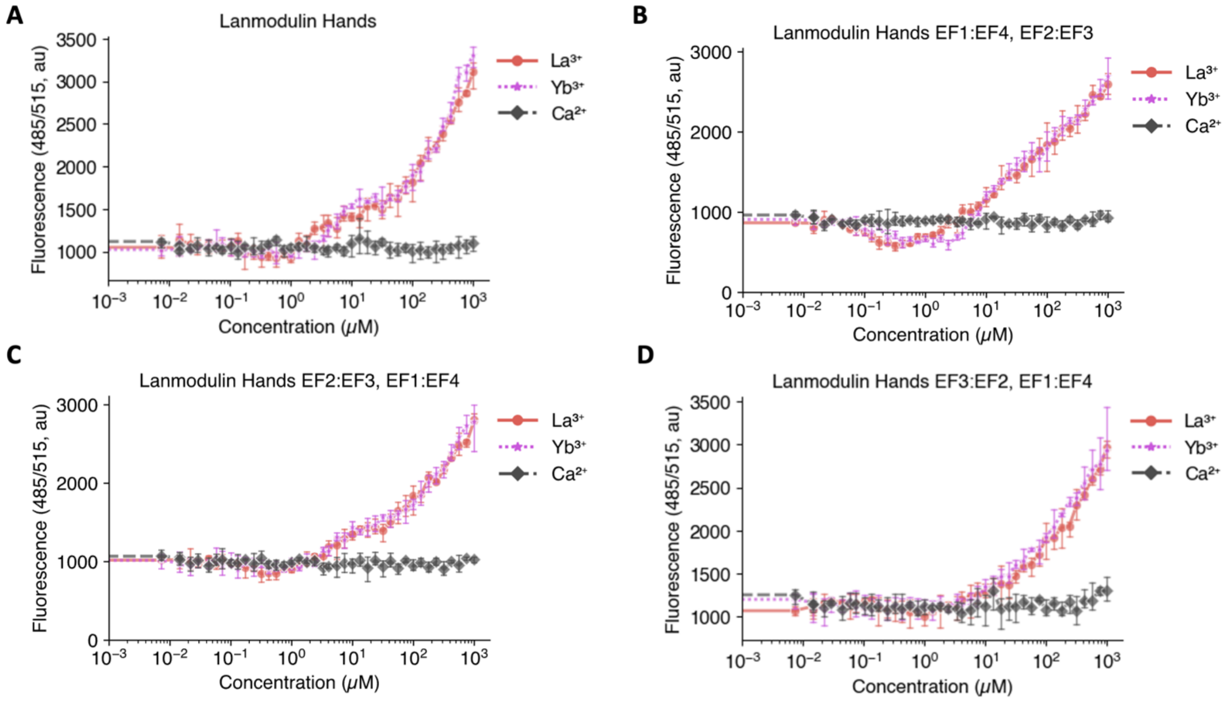

**Supplemental Figure 1: Permutation of Lanmodulin hand order has minimal impact on responsiveness of the sensor**

Plots depict the fluorescence measurements of 4 lanmodulin-binding GCamP variants containing *mex*-LanM EF hands in different N-to-C orders. 500nM of protein was incubated with varying concentrations of the native GCaMP ligand calcium, and lightest (lanthanum) and heaviest (ytterbium) f block lanthanides, then fluorescence measured. Points and Error bars represent the mean and standard deviation of three technical replicates from the same protein purification and working dilution. Graphs of two additional protein purifications can be found in supplemental figures 3 and 5. In **A**, the LanM EF hands are placed in the N-to-C order that they occur in *mex*-LanM sequence (data are identical to Fig. 1C and included here for reference). In **B,C,D** the N-to-C hand order is swapped so that each set of paired hands in one lobe of calmodulin contains the first (EF1) and forth (EF4) EF hands and the other lobe of calmodulin contains the second (EF2) and third (EF3) EF hands. These arrangements more closely mimic the arrangement of the four EF hands in native *mex*-LanM.[4] Specifically, in **B**, the N-to-C order of the EF hands is EF1, EF4, EF2, EF3. In **C**, the N-to-C order of the EF hands is EF2, EF3, EF1, EF4. In **D**, the N-to-C order of the EF hands is EF3, EF2, EF1, EF4.

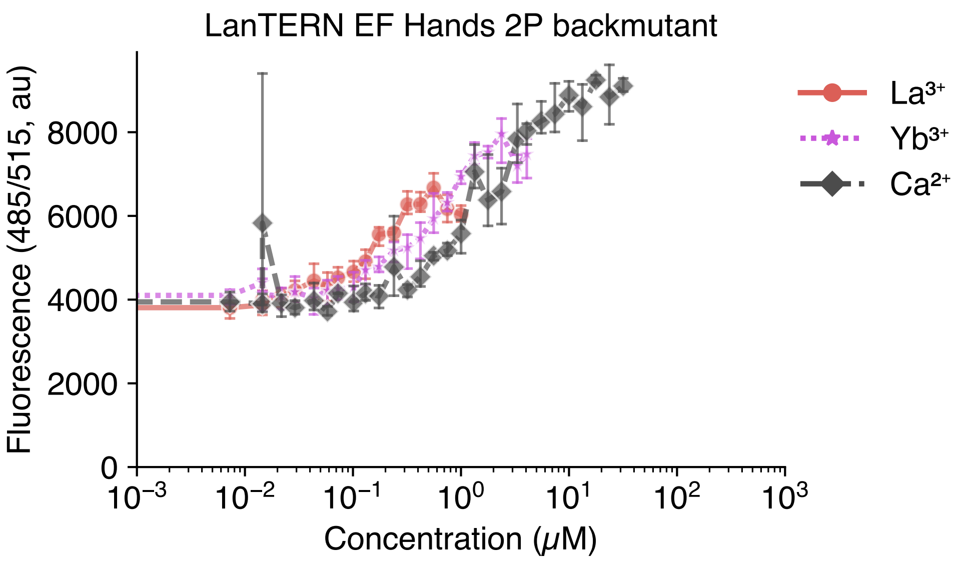

**Supplemental Figure 2: Removal of second position proline restores calcium-dependent activity and lowers lanthanide-dependent activity**

Plots depict the fluorescence measurements of a LanTERN variant where the 2nd position proline of each EF hand has been back mutated to the calmodulin sequence (see Fig. 1A). 500nM of protein was incubated with varying concentrations of the native GCaMP ligand calcium, and lightest (lanthanum) and heaviest (ytterbium) f block lanthanides, then fluorescence measured. Points and Error bars represent the mean and standard deviation of three technical replicates from the same protein purification and working dilution. Lines represent a linear interpolation between points. Graphs of two additional protein purifications can be found in Supplemental Figure 7.

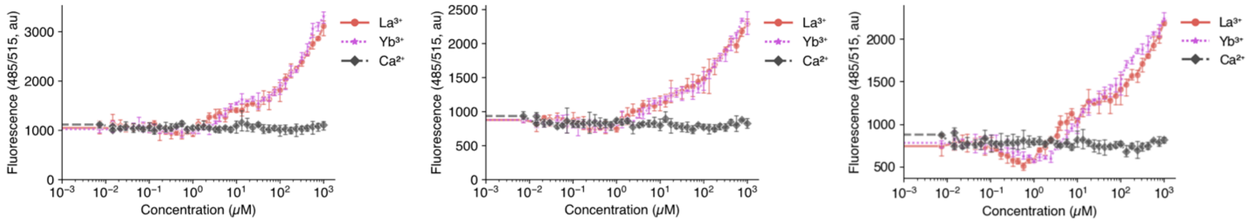

**Supplemental Figure 3: Independent replicates of Fig. 1C (LanM-GCaMP).**

Plots depict the fluorescence measurements of three independent protein purifications of LanM-GCaMP. 500nM of protein was incubated with varying concentrations of the native GCaMP ligand calcium, and lightest (lanthanum) and heaviest (ytterbium) f block lanthanides, then fluorescence measured. Points and Error bars represent the mean and standard deviation of three technical replicates from the same protein purification and working dilution.

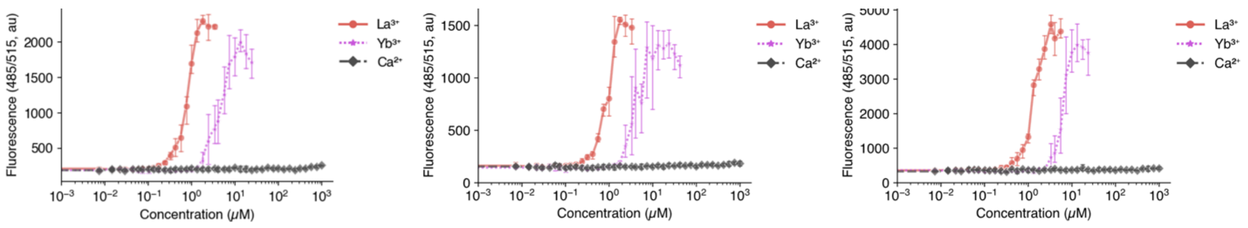

**Supplemental Figure 4: Independent replicates of Fig. 1D (LanTERN)**

Plots depict the fluorescence measurements of three independent protein purifications of LanTERN. 500nM of protein was incubated with varying concentrations of the native GCaMP ligand calcium, and lightest (lanthanum) and heaviest (ytterbium) f block lanthanides, then fluorescence measured. Points and Error bars represent the mean and standard deviation of three technical replicates from the same protein purification and working dilution.

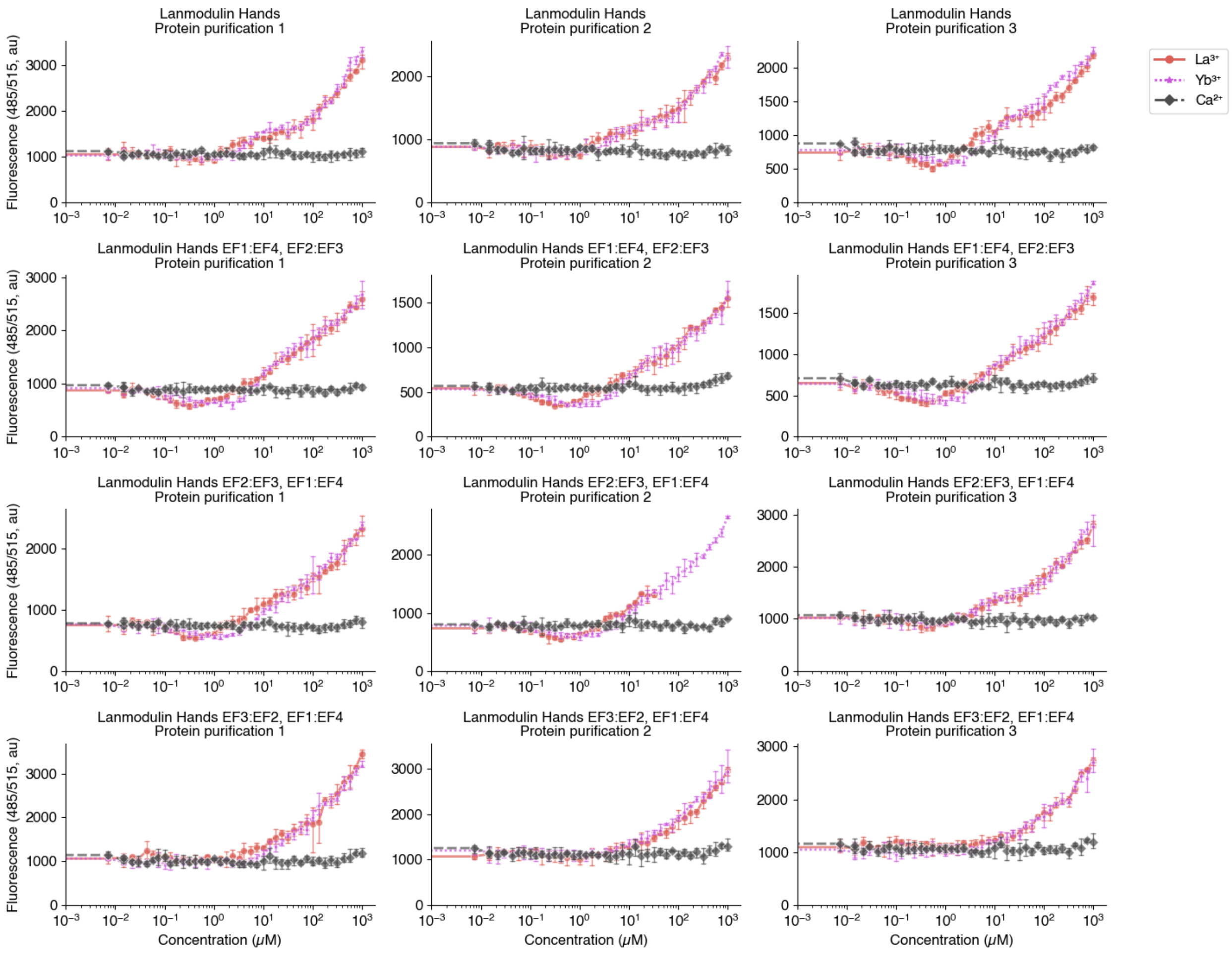

**Supplemental Figure 5: Independent replicates of Fig. S1. (Permutation of Lanmodulin hand order)**

Plots depict the fluorescence measurements of independent protein purifications (protein constructs are labeled in figure panels). 500nM of protein was incubated with varying concentrations of the native GCaMP ligand calcium, and lightest (lanthanum) and heaviest (ytterbium) f block lanthanides, then fluorescence measured. Points and Error bars represent the mean and standard deviation of three technical replicates from the same protein purification and working dilution. Rows represent constructs (see Fig. S1 for details on naming), and columns protein purifications.

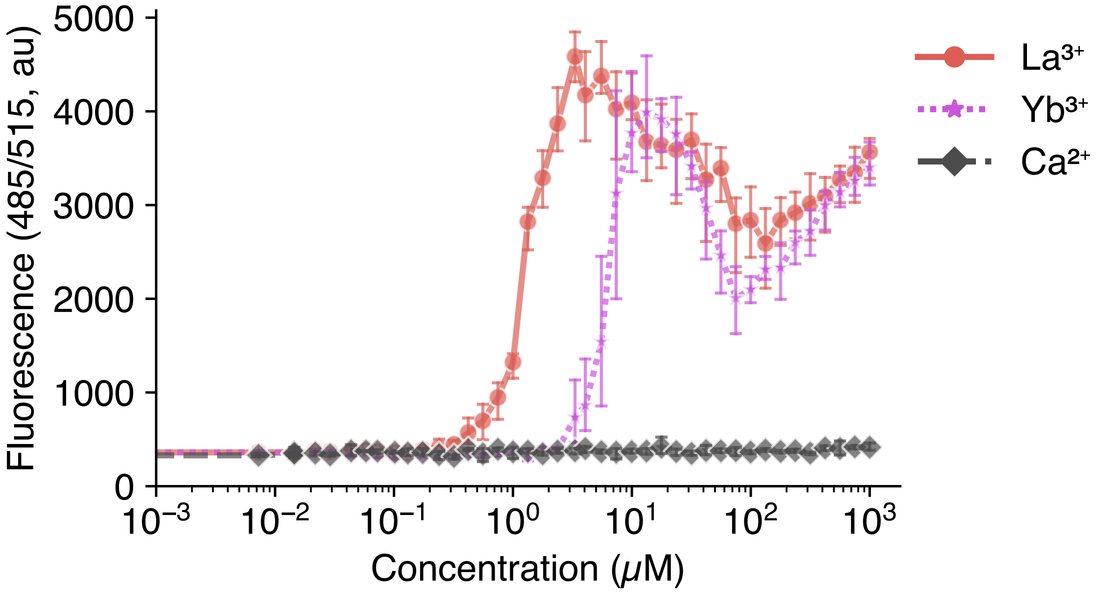

**Supplemental Figure 6: LanTERN dynamics above $Ln_{max}$.**

Plots depict the fluorescence measurements of 500nM of LanTERN in the presence of different concentrations of calcium, and lightest (lanthanum) and heaviest (ytterbium) f block lanthanides, including those above the working range of the sensor. Points and Error bars represent the mean and standard deviation of three technical replicates from the same protein purification and working dilution. Lines represent a linear interpolation between points. The protein purification shown is the same as in Fig. 1B.

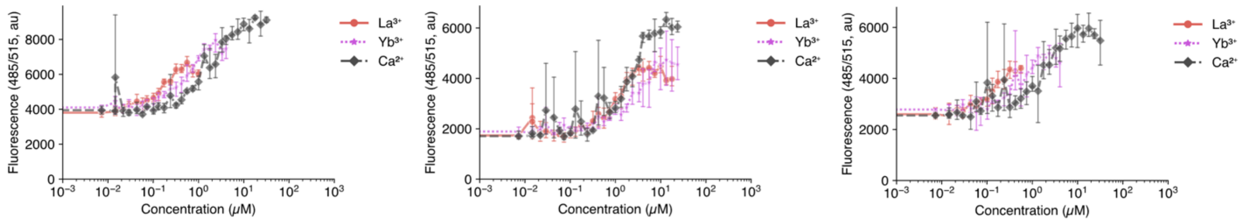

**Supplemental Figure 7: Independent replicates of Fig. S2 (Proline backmutants in LanTERN).**

Plots depict the fluorescence measurements of three independent protein purifications of a LanTERN variant where the 2nd position proline of each EF handhas been back mutated to the calmodulin sequence. 500nM of protein was incubated with varying concentrations of the native GCaMP ligand calcium, and lightest (lanthanum) and heaviest (ytterbium) f block lanthanides, then fluorescence measured. Points and Error bars represent the mean and standard deviation of three technical replicates from the same protein purification and working dilution. Lines represent a linear interpolation between points.

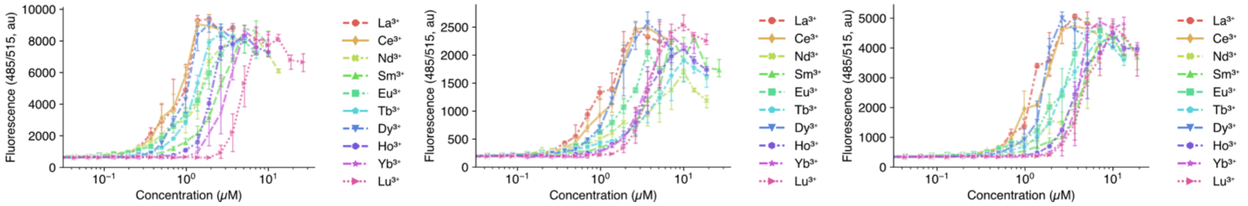

**Supplemental Figure 8: Independent relicates of Fig. 2 (LanTERN response to 10 lanthanides)**

Plots depict the fluorescence measurements of three independent protein purifications of LanTERN. 500nM of protein was incubated with varying concentrations of lanthanides (listed in order of atomic number), then fluorescence measured. Points and Error bars represent the mean and standard deviation of three technical replicates from the same protein purification and working dilution. Lines represent a linear interpolation between points.

### Supplemental Tables

#### Supplemental Table 1: EC50 values of LanTERN

### Supplemental Tables

#### Supplemental Table 1: EC50 values of LanTERN

| | Protein Purification 1 | Protein Purification 2 | Protein Purification 3 | Mean | Standard Deviation |
|---|---|---|---|---|---|
| $La^{3+}$ | 0.93 µM | 0.90 µM | 1.21 µM | 1.01 µM | 0.14 µM |
| $Ce^{3+}$ | 0.80 µM | 1.46 µM | 1.49 µM | 1.25 µM | 0.31 µM |
| $Nd^{3+}$ | 1.38 µM | 3.20 µM | 3.92 µM | 2.83 µM | 1.07 µM |
| $Sm^{3+}$ | 2.33 µM | 3.58 µM | 4.21 µM | 3.37 µM | 0.78 µM |
| $Eu^{3+}$ | 1.32 µM | 1.81 µM | 2.40 µM | 1.84 µM | 0.44 µM |
| $Tb^{3+}$ | 1.20 µM | 3.59 µM | 2.39 µM | 2.39 µM | 0.98 µM |
| $Dy^{3+}$ | 1.01 µM | 1.44 µM | 1.58 µM | 1.34 µM | 0.24 µM |
| $Ho^{3+}$ | 1.92 µM | 3.11 µM | 3.51 µM | 2.85 µM | 0.67 µM |
| $Yb^{3+}$ | 2.97 µM | 3.09 µM | 3.99 µM | 3.35 µM | 0.45 µM |
| $Lu^{3+}$ | 4.54 µM | 3.58 µM | 4.50 µM | 4.21 µM | 0.44 µM |

*EC50 values from experiments in Fig. 2 and Fig. S7. See Supplemental Methods and included Python notebooks for details on EC50 calculation. Mean and standard deviation are reported in the main text of Figure 2B.*

#### Supplemental Table 2: Oligos used in this study

| ID | Purpose | Sequence |
|---|---|---|
| EMJ_P088 | Amplification of GcaMP vector | gttatattccagtttatgccccag |
| EMJ_P089 | Amplification of GcaMP vector | GCGACTCTAGATCATAATCAGCC |
| oEMJ_1200 | Calmodulin for LanTERN (pEMJ_042) | See supplementary sequence file |
| oEMJ_1201 | Calmodulin with native LanM hands (pEMJ_155) | See supplementary sequence file |
| oEMJ_1202 | Calmodulin with LanTERN hands and 2P back mutation (pEMJ_116) | See supplementary sequence file |
| oEMJ_1203 | Calmodulin with native LanM hands; permuted EF2:EF3, EF1:EF4 (pEMJ_156) | See supplementary sequence file |
| oEMJ_1204 | Calmodulin with native LanM hands; permuted EF1:EF4, EF2:EF3 (pEMJ_157) | See supplementary sequence file |

| | | |
|---|---|---|
| oEMJ_1205 | Calmodulin with native LanM hands; permuted EF3:EF2, EF1:EF4 (pEMJ_158) | See supplementary sequence file |

##### Supplemental Table 3: Constructs Used

| ID | Name | Figures |
|---|---|---|
| pEMJ_042 | pRSET LanTERN | Figure 1d, Figure 2, Supplemental figure 4, Supplemental figure 7 |
| pEMJ_116 | pRSET LanTERN 2P backmutant | Supplemental figures 2, 6 |
| pEMJ_155 | pRSET GCaMP native LanM hands | Figure 1c, Supplemental figure 3 |
| pEMJ_156 | pRSET GCaMP native LanM hands; permuted EF2:EF3, EF1:EF4 | Supplemental figure 2, Supplemental figure 5 |
| pEMJ_157 | pRSET GCaMP native LanM hands; permuted EF1:EF4, EF2:EF3 | Supplemental figure 2, Supplemental figure 5 |
| pEMJ_158 | pRSET GCaMP native LanM hands; permuted EF3:EF2, EF1:EF4 | Supplemental figure 2, Supplemental figure 5 |

### Catalog numbers of materials appearing in methods

- NEBuilder® HiFi DNA Assembly Master Mix (NEB: E2621)
- Q5® High-Fidelity 2X Master Mix (NEB: M0492)
- DpnI (NEB: R0176)
- DNA Clean & Concentrator-5 (Zymo: D4013)
- NEB® 5-alpha Competent E. coli (High Efficiency) (NEB: C2987)
- EconoSpin Mini Spin Column(Epoch Life Sciences: 1920-050)
- NiCo21(DE3) *E. coli* (NEB: C2529)
- Falcon® 50 mL High Clarity PP Centrifuge Tube, Conical Bottom, Sterile (Corning: 352098)
- Pierce™ Disposable Columns, 10 mL (ThermoFisher: 29924)
- Chitin Resin (S6651)
- His60 Ni Superflow Resin (Takara: 635661)
- Amicon® Ultra-15 Centrifugal Filter Unit (Millipore: UFC901024)
- Econo-Pac 10DG Desalting Column (Bio-Rad: 7322010)
- Pierce™ BCA Protein Assay (ThermoFisher: 23225)

- MICROPLATE, 384 WELL, PP, F-BOTTOM (Greiner Bio-One: 781209)
- Lanthanide Supplies
    - Lanthanum(III) nitrate hexahydrate (Sigma Aldrich: 331937)
    - Cerium(III) nitrate hexahydrate, REacton®, 99.99% (REO), (Alpha Aesar: 11330)
    - Neodymium(III) nitrate hexahydrate (Sigma Aldrich: 289175)
    - Samarium(III) nitrate hexahydrate, 99.999% trace metals basis (Sigma Aldrich: 518247)
    - Europium(III) nitrate hydrate (Sigma Aldrich: 254061)
    - Terbium(III) nitrate hexahydrate (Sigma Aldritch: 217212)
    - Dysprosium(III) nitrate pentahydrate, REacton™, 99.99% (REO), (ThermoFisher: 011315.14)
    - Holmium(III) nitrate pentahydrate, 99.99% trace metals basis (Sigma Aldrich: 229687)
    - Ytterbium(III) nitrate pentahydrate, 99.999% (Sigma Aldritch: 217220)
    - Lutetium(III) nitrate hydrate, REacton™, 99.99% (ThermoFisher: 011258.04)

### Supplemental Data Information

The supplemental data attachment includes the raw and processed data from all experiments and the Python notebooks used to process the data. It also includes annotated GenBank files of all sequences reported in the study.

The supplemental data occasionally includes constructs and protein purifications not mentioned in the text (for instance, a BCA may include a failed protein purification); these were maintained to avoid altering the raw data.

Data refers to plasmids using the same notation as in the text. Protein purifications are labeled using the notation EMJ_RXXX. Some plate reader experiments contain additional data from constructs not referred to in the manuscript; when this is the case, that data is excluded in the processing code in "prepare_data.ipynb" and explained with a comment. For transparency, it was decided that it is preferable to avoid altering any of the raw data. While the Excel exports of the raw data for 3 of the 4 BCAs used to determine the protein concentration are present, the original plate reader files were unfortunately not maintained. The authors regret this omission. To analyze the plate reader data using the raw Excel outputs of the instruments, simply run the code in the notebook present in each plate reader data folder. Then run prepare_data.ipynb and calculate_half_max.ipynb. Versions of the plots found in the text can be generated using their associated .ipynb files.

### References

(1) Studier, F. W. Stable Expression Clones and Auto-Induction for Protein Production in E. Coli. In *Structural Genomics: General Applications*; Chen, Y. W., Ed.; Methods in Molecular

Biology; Humana Press: Totowa, NJ, 2014; pp 17–32. https://doi.org/10.1007/978-1-62703-691-7_2.

(2) Mattocks, J. A.; Ho, J. V.; Cotruvo, J. A. A Selective, Protein-Based Fluorescent Sensor with Picomolar Affinity for Rare Earth Elements. *J. Am. Chem. Soc.* **2019**, *141* (7), 2857–2861. https://doi.org/10.1021/jacs.8b12155.

(3) Sali, A.; Blundell, T. L. Comparative Protein Modelling by Satisfaction of Spatial Restraints. *J. Mol. Biol.* **1993**, *234* (3), 779–815. https://doi.org/10.1006/jmbi.1993.1626.

(4) Cook, E. C.; Featherston, E. R.; Showalter, S. A.; Cotruvo, J. A. Structural Basis for Rare Earth Element Recognition by Methylobacterium Extorquens Lanmodulin. *Biochemistry* **2019**, *58* (2), 120–125. https://doi.org/10.1021/acs.biochem.8b01019.
