## Supplemental Information 2 for "LanTERN: a fluorescent sensor that specifically binds lanthanides": calculate_half_max.ipynb

```
# %% [code]
import pandas as pd
import matplotlib.pyplot as plt
import seaborn as sns
import numpy as np
from sklearn.linear_model import LinearRegression
import os
import scipy
sns.set_palette(sns.color_palette('colorblind'))

# %% [code]
def q(x):
    order = ['La','Ce','Nd','Sm','Eu','Tb','Dy','Ho','Yb','Lu']
    ion_symbol=f'\N{SUPERSCRIPT THREE}\N{SUPERSCRIPT PLUS SIGN}'
    order = [i +ion_symbol for i in order]
    d =dict(zip(order,[i for i in range(len(order))]))
    return x.map(d)


def calculate_half_max(row):
    row = row.sort_values('Concentration (µM)')
    max_val = row['485/515'].max()
    last_concentration = None
    last_value = None
    for index, val in row.iterrows():
        
        if val['485/515']/max_val >0.5:
            x= np.array([last_concentration,val['Concentration (µM)']]).reshape(-1,1)
            y= np.array([last_value/max_val,val['485/515']/max_val]).reshape(-1,1)
            return LinearRegression().fit(y,x).predict([[0.5]])[0][0]
            
            
        last_concentration = val['Concentration (µM)']
        last_value =val['485/515']
df=pd.read_csv('fig1_data.csv')        

#EC50s not calculated for other constructs as they do not reach a firm steady state
x=df[(df['Ion']!='Ca²⁺')&(df['Construct']=='LanTERN EF Hands')].groupby(['Protein Prep','Ion','Concentration (µM)']).mean().reset_index()

x.groupby(['Protein Prep','Ion']).apply(calculate_half_max).reset_index().sort_values('Protein Prep').sort_values('Ion',key=q).T.to_excel('fig1_EC50_calculations.xlsx',index=False)


# %% [code]
df=pd.read_csv('fig2_data.csv') 


def q(x):
    order = ['La','Ce','Nd','Sm','Eu','Tb','Dy','Ho','Yb','Lu']
    ion_symbol=f'\N{SUPERSCRIPT THREE}\N{SUPERSCRIPT PLUS SIGN}'
    order = [i +ion_symbol for i in order]
    d =dict(zip(order,[i for i in range(len(order))]))
    d['R027']=1
    d['R039']=2
    d['R040']=3
    d['R041']=4
    return x.map(d)


x=df[(df['Ion']!='Ca²⁺')&(df['Construct']=='LanTERN EF Hands')].groupby(['Protein Prep','Ion','Concentration (µM)']).mean().reset_index()
x.groupby(['Protein Prep','Ion']).apply(calculate_half_max).reset_index().sort_values('Protein Prep').sort_values(['Ion','Protein Prep'],key=q).T.to_excel('fig2_EC50_calculations.xlsx',index=False)


# %% [code]
```
