## Supplemental Information 2 for "LanTERN: a fluorescent sensor that specifically binds lanthanides": Figure 1.ipynb

```
# %% [code]
import pandas as pd
import matplotlib.pyplot as plt
import seaborn as sns
import numpy as np
import matplotlib as mpl
from matplotlib.ticker import (MultipleLocator, AutoMinorLocator)
import matplotlib.ticker as ticker
from matplotlib.ticker import LogLocator, LogFormatter,LogFormatterMathtext
import os
mpl.rcParams['font.family']='Helvetica'

rc = {'scatter.markersize' : 2.5,'lines.linewidth':2,'font.size':12,'axes.titlesize':12}
error_bar_capsize=2
alpha = 0.7
elinewidth=1.0

error_kws={'capsize':error_bar_capsize,'elinewidth':elinewidth}#,'alpha':alpha}

color_map_name= 'hls'
figsize=(4.36,4.36)

# %% [code]
main_data=pd.read_csv('fig1_data.csv')
construct_key=dict(set(zip(main_data['Protein Prep'].values.tolist(),main_data['Construct'].values.tolist())))

# %% [code]
figure_number= 'fig1'

# %% [code]
def get_marker_key():
    order = ['La','Ce','Nd','Sm','Eu','Tb','Dy','Ho','Yb','Lu']
    ion_symbol=f'\N{SUPERSCRIPT THREE}\N{SUPERSCRIPT PLUS SIGN}'
    order = [i +ion_symbol for i in order]+['Ca'+f'\N{SUPERSCRIPT Two}\N{SUPERSCRIPT PLUS SIGN}']
    return dict(zip(order,['o', 'd','X',  '^', 's', 'p', 'v', 'h', '*','>','D']))#,["o", "^","h", "P",'d','p','x','v','*','+','>']))
markers=get_marker_key()


# %% [code]
order = ['La','Yb']
ion_symbol=f'\N{SUPERSCRIPT THREE}\N{SUPERSCRIPT PLUS SIGN}'
order = [i +ion_symbol for i in order] +[f'Ca\N{SUPERSCRIPT TWO}\N{SUPERSCRIPT PLUS SIGN}'] 
color_map_name= 'hls'
color_palette = sns.color_palette(color_map_name, n_colors=main_data['Ion'].nunique())

color_palette = {order[0]:(0.86, 0.3712, 0.33999999999999997),
                order[1]:(0.7871999999999999, 0.33999999999999997, 0.86),
                 order[2]: (0.3,0.3,0.3)}

# %% [code]
if not os.path.exists(figure_number):
    os.mkdir(figure_number)

# %% [code]
for prep in main_data['Protein Prep'].unique():
    construct = construct_key[prep]
    construct=construct.replace('LanTERN EF Hands w/o proline','LanTERN EF Hands 2P backmutant')
    if not os.path.exists(os.path.join(figure_number,construct)):
        os.mkdir(os.path.join(figure_number,construct))
    fp = os.path.join(figure_number,construct,f'{prep}.svg')
    
    with sns.plotting_context(rc=rc):
        fig,ax= plt.subplots(1,1,figsize=figsize)
        df=main_data[main_data['Protein Prep']==prep]
        sns.lineplot(data=df,
                     x='Concentration (µM)',
                     y='485/515',hue='Ion',
                     ax=ax,
                     palette=color_palette,
                     alpha=alpha,
                     err_kws=error_kws,
                     err_style='bars',
                     hue_order=order,
                     style='Ion',
                     markers=markers)
        ax.set_xscale('log')
        ax.set_ylim(bottom=0)

        # Set major ticks
        major_ticks = np.logspace(-3, 3, num=7) # Gives an array [1e-03, 1e-02, 1e-01, 1e+00, 1e+01, 1e+02, 1e+03]
        ax.set_xticks(major_ticks)
        # Set minor ticks
        minor_locator = LogLocator(subs=np.arange(2, 10) * .1, numticks=12)
        ax.get_xaxis().set_minor_locator(minor_locator)
        ax.get_xaxis().set_minor_formatter(LogFormatter())
        ax.spines['top'].set_visible(False)
        ax.spines['right'].set_visible(False)
        plt.title(construct.replace('LanM','Lanmodulin'))
        plt.legend(frameon=False,loc='upper right',bbox_to_anchor=(1.3,1))
        ax.set_ylabel('Fluorescence (485/515, au)')
        plt.savefig(fp,bbox_inches='tight')
        plt.show()
```
