## Supplemental Information 2 for "LanTERN: a fluorescent sensor that specifically binds lanthanides": 8-2-2023.ipynb

```
# %% [code]
import pandas as pd
import matplotlib.pyplot as plt
import seaborn as sns
import numpy as np
sns.set_palette(sns.color_palette('colorblind'))

# %% [code]
key=pd.read_excel('8-2-conc-plateset-1.xlsx')
key=key.dropna(subset='Single fluid concentration')
key=key[['Non-random well','Well contents','Single fluid concentration']]
key['Well contents']=key['Well contents'].str.replace(' - 1:3','')
key

# %% [code]
key2=pd.read_excel('8-2-conc-plateset-2.xlsx')
key2=key2.dropna(subset='Single fluid concentration')
key2=key2[['Non-random well','Well contents','Single fluid concentration']]
key2['Well contents']=key2['Well contents'].str.replace(' - 1:3','')
key2

# %% [code]
def load_data(excel_filename,sheet_name,key):
    def apply_pad(string):
        if string in [str(i) for i in range(10)]:
            return '0'+string
        return string

    df=pd.read_excel(excel_filename,sheet_name=sheet_name,skiprows=27,usecols ='B:Z').melt('Unnamed: 1')
    df=df.rename(columns={'Unnamed: 1':'Row','variable':'Column','value':'485/515'})
    df['Column']=df['Column'].astype(str).apply(apply_pad)
    df['Well']=df['Row']+df['Column'].astype(str)
    df=df.dropna()
    df=df.merge(key,how='left',left_on='Well',right_on='Non-random well')

    for i in df['Well contents'].unique():
        if type(i) is str:
            df.loc[df[df['Well contents'].isna()].index[0:3],'Well contents']=i

    df['Single fluid concentration']=df['Single fluid concentration'].fillna(0)
    df=df.dropna(subset='Well contents')
    df['replicate']=1
    df.loc[df.duplicated(subset=['Well contents','Single fluid concentration','replicate']),'replicate']+=1
    df.loc[df.duplicated(subset=['Well contents','Single fluid concentration','replicate']),'replicate']+=1
    df=df.rename(columns={'Well contents':'Ion','Single fluid concentration':'Concentration (µM)'}).drop('Non-random well',axis=1)

    def functional_lowest(x):
        return df[(df['Ion']==x['Ion'])&(df['replicate']==x['replicate'])]['Concentration (µM)'].idxmin()

    df['min_index']=df.apply(functional_lowest,axis=1)


    def check(row):
            #safety check
            check_row=df.loc[row['min_index']]
            if check_row['Ion']!= row['Ion'] or check_row['Concentration (µM)']!= 0:
                return False
            else:
                return True
    if not (len(df.apply(check,axis=1).unique())==1 and True in df.apply(check,axis=1).unique()):
        raise Exception('Error Normalization Incorrect')

    def prepare_normalizer(min_index):
        return df.loc[min_index]['485/515']
    df['Normalizer 485/515']=df['min_index'].apply(prepare_normalizer)
    df['Fold Change (485/515)']=df['485/515']/df['Normalizer 485/515']
    df['Log2 Fold Change (485/515)']=np.log2(df['Fold Change (485/515)'])
    df['Protein Prep']=sheet_name
    return df

# %% [code]
excel_filename='2023-08-02-plates-1.xlsx'
sheets =['R027','R040','R041']
df=pd.concat([load_data(excel_filename,sheet_name,key) for sheet_name in sheets],ignore_index=True)
excel_filename='2023-08-02-plates-2.xlsx'
df=pd.concat([load_data(excel_filename,sheet_name,key2) for sheet_name in sheets] +[df],ignore_index=True)

plasmid_keys = {'R027':'pEMJ_042','R040':'pEMJ_042','R041':'pEMJ_042'}
name_keys = {'pEMJ_060':'LanM Hands','pEMJ_062':'LanM Hands EF2:EF3, EF1:EF4','pEMJ_042':'LanTERN EF Hands'}
df['plasmid']=df['Protein Prep'].map(plasmid_keys)
df['Construct']=df['plasmid'].map(name_keys)
df['Experiment Date']= '8-2-2023'
df.to_csv('08-02-2023_processed.csv',index=False)
```
