## Supplemental Information 2 for "LanTERN: a fluorescent sensor that specifically binds lanthanides": Prepare data.ipynb

```
# %% [code]
import pandas as pd
import matplotlib.pyplot as plt
import numpy as np
import os
import scipy

# %% [markdown]
### Fig1

# %% [code]
df=pd.concat([pd.read_csv('Plate_Reader_Data/2023-07-26/7-26-2023_processed.csv'),
              pd.read_csv('Plate_Reader_Data/2023-07-27/7-27-2023_processed.csv'),
            pd.read_csv('Plate_Reader_Data/2023-09-11/9-11-2023_processed.csv')],ignore_index=True)
col_order=['LanTERN EF Hands',
 'LanM Hands',
'LanM Hands EF2:EF3, EF1:EF4',
'LanM Hands EF1:EF4, EF2:EF3',
'LanM Hands EF3:EF2, EF1:EF4','LanTERN EF Hands w/o proline']


order = ['La','Ce','Nd','Sm','Eu','Tb','Ho','Yb','Lu']
ion_symbol=f'\N{SUPERSCRIPT THREE}\N{SUPERSCRIPT PLUS SIGN}'
order = [i +ion_symbol for i in order] +[f'Ca\N{SUPERSCRIPT TWO}\N{SUPERSCRIPT PLUS SIGN}'] 
order_og  = ['La','Ce','Nd','Sm','Eu','Tb','Ho','Yb','Lu','Ca']
for i in range(len(order)):
     df.loc[:,'Ion']=df['Ion'].str.replace(order_og[i],order[i])
        

df['style']='True'

def q(x):
    d =dict(zip(order,[i for i in range(len(order))]))
    return x.map(d)

# %% [code]
#This step excludes some of the plasmids. In particular it excludes
#pEMJ_060-pEMJ_064. The reason for this is because during the preparation
#of this manuscript, it was noticed that our wild type LanM hand sequence
#was subtly incorrect, and contained a L->I mutation in the 8th position
#of the second EF hand of the above constructs. Because data from some of
#the other constructs was collected during the same experiment, we chose
#to maintain this data's presence so as to avoid modifying the original data
#files. We then built and tested corrected constructs (pEMJ_155-pEMJ_158),
#which are the ones used in the figures. Notably there is little to no 
#difference in the behavior of the EF2 L8I mutant, and the wild type hands 
#which is unsurprising. 

excluded_plasmids =['pEMJ_060','pEMJ_062','pEMJ_063','pEMJ_064']
df=df[~(df['plasmid'].isin(excluded_plasmids))]
df.plasmid.unique()

# %% [code]
#Filter titration data until maximum value is reach + 3 extra points 
#This is performed because after reaching the maximum value the relationship
#between concentration and fluorescence becomes slightly unusual and going down 
#as concentrations go up and then increasing. Similar findings seem to occur in
#the titration curve of Cortruvo et al. 2018 and Mattocks et al. 2019 

num_extra = 3
out = {}
tester= df.copy()
tester=tester.groupby(['Protein Prep','Ion','Concentration (µM)']).mean().reset_index()
tester['rise started']=False
tester.loc[tester['Fold Change (485/515)']>=1.5,'rise started']=True


for prep in tester['Protein Prep'].unique():
    out[prep]={}
    for ion in tester.Ion.unique():
        data = tester[(tester['Protein Prep']==prep)&(tester['Ion']==ion)&(tester['rise started']==True)]
        if len(data) ==0:
                out[prep][ion]=tester[(tester['Protein Prep']==prep)&(tester['Ion']==ion)]['Concentration (µM)'].max()
        else:
            l=data.sort_values('Concentration (µM)',ascending=True)['Concentration (µM)'].values.tolist()
            

            flx =data.sort_values('Concentration (µM)',ascending=True)['Fold Change (485/515)'].values.tolist()

            highest_value=flx[0]
            counter = 0
            for i in flx[1:]:
                if counter ==num_extra:
                    break
                if i>highest_value:
                    highest_value=i
                    counter=0
                else:
                    counter+=1

            for i in l[flx.index(highest_value):flx.index(highest_value)+num_extra]:
                pass
                out[prep][ion]=i
plotting_df=df.copy()
for prep,d in out.items():
    for ion,value in d.items():
        plotting_df=plotting_df[~((plotting_df['Protein Prep']==prep)&(plotting_df['Ion']==ion)&(plotting_df['Concentration (µM)']>value))]

# %% [code]
plotting_df.to_csv('fig1_data.csv',index=False)

# %% [markdown]
### Fig 2

# %% [code]
df=pd.concat([pd.read_csv('Plate_Reader_Data/2023-08-02/08-02-2023_processed.csv'),
              pd.read_csv('Plate_Reader_Data/2023-08-07/08-07-2023_processed.csv')],ignore_index=True)


#The protein prep R040 seemed to behave poorly and on inspection appears to have been partially degraded
#due to improper storage. An older protein prep R027 with known good behavior was inspected and used in its
#place for figure 2. The existing R040 data was maintained for the sake of transparency and 
#providing unaltered data files
df=df[df['Protein Prep']!='R040']

order = ['La','Ce','Nd','Sm','Eu','Tb','Dy','Ho','Yb','Lu']
ion_symbol=f'\N{SUPERSCRIPT THREE}\N{SUPERSCRIPT PLUS SIGN}'
order = [i +ion_symbol for i in order] +[f'Ca\N{SUPERSCRIPT TWO}\N{SUPERSCRIPT PLUS SIGN}'] 
order_og  = ['La','Ce','Nd','Sm','Eu','Tb','Dy','Ho','Yb','Lu','Ca']
for i in range(len(order)):
     df.loc[:,'Ion']=df['Ion'].str.replace(order_og[i],order[i])
        

df['style']='True'


# %% [code]
#Filter titration data until maximum value is reach + 3 extra points 
num_extra = 3
out = {}
tester= df.copy()
tester=tester.groupby(['Protein Prep','Ion','Concentration (µM)']).mean().reset_index()
tester['rise started']=False
tester.loc[tester['Fold Change (485/515)']>=1.5,'rise started']=True


for prep in tester['Protein Prep'].unique():
    out[prep]={}
    for ion in tester.Ion.unique():
        data = tester[(tester['Protein Prep']==prep)&(tester['Ion']==ion)&(tester['rise started']==True)]
        if len(data) ==0:
                out[prep][ion]=tester[(tester['Protein Prep']==prep)&(tester['Ion']==ion)]['Concentration (µM)'].max()
        else:
            l=data.sort_values('Concentration (µM)',ascending=True)['Concentration (µM)'].values.tolist()
            

            flx =data.sort_values('Concentration (µM)',ascending=True)['Fold Change (485/515)'].values.tolist()

            highest_value=flx[0]
            counter = 0
            for i in flx[1:]:
                if counter ==num_extra:
                    break
                if i>highest_value:
                    highest_value=i
                    counter=0
                else:
                    counter+=1

        for i in l[flx.index(highest_value):flx.index(highest_value)+num_extra]:
            pass
            out[prep][ion]=i
plotting_df=df.copy()
for prep,d in out.items():
    for ion,value in d.items():
        plotting_df=plotting_df[~((plotting_df['Protein Prep']==prep)&(plotting_df['Ion']==ion)&(plotting_df['Concentration (µM)']>value))]

# %% [code]
plotting_df.to_csv('fig2_data.csv',index=False)
```
