## Supplemental Information 2 for "LanTERN: a fluorescent sensor that specifically binds lanthanides": Supplemental Figure 5.ipynb

```
# %% [code]
import pandas as pd
import matplotlib.pyplot as plt
import seaborn as sns
import numpy as np
import matplotlib as mpl
from matplotlib.ticker import (MultipleLocator, AutoMinorLocator)
import matplotlib.ticker as ticker
from matplotlib.ticker import LogLocator, LogFormatter,LogFormatterMathtext
import os
mpl.rcParams['font.family']='Helvetica'

rc = {'scatter.markersize' : 2.5,'lines.linewidth':2,'font.size':12,'axes.titlesize':12}
error_bar_capsize=2
alpha = 0.7
elinewidth=1.0

error_kws={'capsize':error_bar_capsize,'elinewidth':elinewidth}#,'alpha':alpha}

color_map_name= 'hls'
figsize=(4.36,2.81)

# %% [code]
main_data=pd.read_csv('fig1_data.csv')
construct_key=dict(set(zip(main_data['Protein Prep'].values.tolist(),main_data['Construct'].values.tolist())))
main_data=main_data[~(main_data['Construct'].isin(['LanTERN EF Hands','LanTERN EF Hands w/o proline']))]

# %% [code]
def get_marker_key():
    order = ['La','Ce','Nd','Sm','Eu','Tb','Dy','Ho','Yb','Lu']
    ion_symbol=f'\N{SUPERSCRIPT THREE}\N{SUPERSCRIPT PLUS SIGN}'
    order = [i +ion_symbol for i in order]+['Ca'+f'\N{SUPERSCRIPT Two}\N{SUPERSCRIPT PLUS SIGN}']
    return dict(zip(order,['o', 'd','X',  '^', 's', 'p', 'v', 'h', '*','>','D']))#,["o", "^","h", "P",'d','p','x','v','*','+','>']))
markers=get_marker_key()


# %% [code]
order = ['La','Yb']
ion_symbol=f'\N{SUPERSCRIPT THREE}\N{SUPERSCRIPT PLUS SIGN}'
order = [i +ion_symbol for i in order] +[f'Ca\N{SUPERSCRIPT TWO}\N{SUPERSCRIPT PLUS SIGN}'] 
color_map_name= 'hls'
color_palette = sns.color_palette(color_map_name, n_colors=main_data['Ion'].nunique())

color_palette = {order[0]:(0.86, 0.3712, 0.33999999999999997),
                order[1]:(0.7871999999999999, 0.33999999999999997, 0.86),
                 order[2]: (0.3,0.3,0.3)}

# %% [code]
for construct in main_data['Construct'].unique():
    for i, protein_prep in enumerate(main_data.loc[main_data['Construct']==construct,'Protein Prep'].unique()):
        main_data['Protein Prep']=main_data['Protein Prep'].str.replace(protein_prep,f'Protein purification {i+1}')

# %% [code]
main_data=main_data.rename(columns={'Protein Prep':'Protein purification'})

# %% [code]
constructs=['LanM Hands','LanM Hands EF1:EF4, EF2:EF3', 
            'LanM Hands EF2:EF3, EF1:EF4','LanM Hands EF3:EF2, EF1:EF4']
figsize=(13.08,11.24)

with sns.plotting_context(rc=rc):
    fig,axes= plt.subplots(4,3,figsize=figsize)
    for i, construct in enumerate(constructs):
        for j, prep in enumerate(main_data['Protein purification'].unique()):
            df=main_data[(main_data['Construct']==construct)&(main_data['Protein purification']==prep)]
            ax = axes[i,j]
            sns.lineplot(data=df,
                     x='Concentration (µM)',
                     y='485/515',hue='Ion',
                     ax=ax,
                     palette=color_palette,
                     alpha=alpha,
                     err_kws=error_kws,
                     err_style='bars',
                     hue_order=order,
                     style='Ion',
                     markers=markers)
            ax.set_xscale('log')
            ax.set_ylim(bottom=0)
            # Set major ticks
            major_ticks = np.logspace(-3, 3, num=7) # Gives an array [1e-03, 1e-02, 1e-01, 1e+00, 1e+01, 1e+02, 1e+03]
            ax.set_xticks(major_ticks)
            # Set minor ticks
            minor_locator = LogLocator(subs=np.arange(2, 10) * .1, numticks=12)
            ax.get_xaxis().set_minor_locator(minor_locator)
            ax.get_xaxis().set_minor_formatter(LogFormatter())
            ax.spines['top'].set_visible(False)
            ax.spines['right'].set_visible(False)
            ax.set_title(construct.replace('LanM','Lanmodulin') + f'\n{prep}')
            #plt.legend(frameon=False,loc='upper right')
            if j==0:
                ax.set_ylabel('Fluorescence (485/515, au)')
            else:
                ax.set_ylabel('')
            if i != 3:
                ax.set_xlabel('')
            ax.get_legend().remove()
    lines, labels = ax.get_legend_handles_labels()
    fig.legend(lines, labels,loc='upper right',bbox_to_anchor=(1.10,0.95),frameon=True)
    plt.tight_layout()
    plt.savefig('supplemental_figure5.png',bbox_inches='tight')
    plt.show()

# %% [code]
```
