## Supplemental Information 2 for "LanTERN: a fluorescent sensor that specifically binds lanthanides": plotting_enviroment.yml

```
name: /Users/EthanJ/opt/anaconda3/envs/plotting
channels:
  - numba
  - conda-forge
  - bioconda
  - defaults
dependencies:
  - _tflow_select=2.2.0=eigen
  - abseil-cpp=20211102.0=he9d5cce_0
  - absl-py=1.3.0=py38hecd8cb5_0
  - adjusttext=0.7.3.1=py_1
  - aiohttp=3.8.3=py38h6c40b1e_0
  - aiosignal=1.2.0=pyhd3eb1b0_0
  - anyio=3.5.0=py38hecd8cb5_0
  - appnope=0.1.2=py38hecd8cb5_1001
  - aragorn=1.2.41=ha5712d3_0
  - argon2-cffi=21.3.0=pyhd3eb1b0_0
  - argon2-cffi-bindings=21.2.0=py38hca72f7f_0
  - argtable2=2.13=h1de35cc_1001
  - asttokens=2.0.5=pyhd3eb1b0_0
  - astunparse=1.6.3=py_0
  - async-timeout=4.0.2=py38hecd8cb5_0
  - attrs=22.1.0=py38hecd8cb5_0
  - babel=2.11.0=py38hecd8cb5_0
  - backcall=0.2.0=pyhd3eb1b0_0
  - barrnap=0.9=hdfd78af_4
  - beautifulsoup4=4.11.1=py38hecd8cb5_0
  - bedtools=2.30.0=h0e31d98_3
  - biopython=1.78=py38h9ed2024_0
  - blas=1.0=mkl
  - blast=2.12.0=h0370960_3
  - bleach=4.1.0=pyhd3eb1b0_0
  - blinker=1.4=py38hecd8cb5_0
  - bottleneck=1.3.5=py38h67323c0_0
  - brotli=1.0.9=hca72f7f_7
  - brotli-bin=1.0.9=hca72f7f_7
  - brotlipy=0.7.0=py38h9ed2024_1003
  - bzip2=1.0.8=h1de35cc_0
  - c-ares=1.18.1=hca72f7f_0
  - ca-certificates=2023.01.10=hecd8cb5_0
  - cachetools=4.2.2=pyhd3eb1b0_0
  - certifi=2022.12.7=py38hecd8cb5_0
  - cffi=1.15.1=py38hc55c11b_0
  - charset-normalizer=2.0.4=pyhd3eb1b0_0
  - click=8.0.4=py38hecd8cb5_0
  - cloudpickle=2.0.0=pyhd3eb1b0_0
  - clustalo=1.2.4=h9722bc1_5
  - clustalw=2.1=hcd10b59_7
  - comm=0.1.2=py38hecd8cb5_0
  - cryptography=38.0.4=py38hf6deb26_0
  - curl=7.84.0=hca72f7f_0
  - cycler=0.11.0=pyhd3eb1b0_0
  - debugpy=1.5.1=py38he9d5cce_0
  - decorator=5.1.1=pyhd3eb1b0_0
  - defusedxml=0.7.1=pyhd3eb1b0_0
  - entrez-direct=16.2=h193322a_0
  - entrypoints=0.4=py38hecd8cb5_0
  - executing=0.8.3=pyhd3eb1b0_0
  - expat=2.4.9=he9d5cce_0
  - fairlearn=0.7.0=pyhd8ed1ab_0
  - fftw=3.3.9=h9ed2024_1
  - flatbuffers=2.0.0=h23ab428_0
  - flit-core=3.6.0=pyhd3eb1b0_0
  - fonttools=4.25.0=pyhd3eb1b0_0
  - freetype=2.11.0=hd8bbffd_0
  - frozenlist=1.3.3=py38h6c40b1e_0
  - gast=0.4.0=pyhd3eb1b0_0
  - giflib=5.2.1=h6c40b1e_3
  - google-auth=2.6.0=pyhd3eb1b0_0
  - google-auth-oauthlib=0.4.4=pyhd3eb1b0_0
  - google-pasta=0.2.0=pyhd3eb1b0_0
  - greenlet=2.0.1=py38hcec6c5f_0
  - grpc-cpp=1.46.1=h067a048_0
  - grpcio=1.42.0=py38ha29bfda_0
  - h5py=3.7.0=py38h4a1dd59_0
  - hdf5=1.10.6=h10fe05b_1
  - hmmer=3.3.2=h9722bc1_2
  - hmmlearn=0.2.8=py38hec72209_1
  - icu=58.2=h0a44026_3
  - idna=3.4=py38hecd8cb5_0
  - imbalanced-learn=0.9.1=pyhd8ed1ab_1
  - importlib-metadata=4.11.3=py38hecd8cb5_0
  - importlib_metadata=4.11.3=hd3eb1b0_0
  - importlib_resources=5.2.0=pyhd3eb1b0_1
  - infernal=1.1.4=pl5321ha5712d3_1
  - intel-openmp=2021.4.0=hecd8cb5_3538
  - ipykernel=6.19.2=py38h01d92e1_0
  - ipympl=0.8.7=pyhd3eb1b0_0
  - ipython=8.4.0=py38hecd8cb5_0
  - ipython_genutils=0.2.0=pyhd3eb1b0_1
  - ipywidgets=7.6.5=pyhd3eb1b0_1
  - jedi=0.18.1=py38hecd8cb5_1
  - jinja2=3.1.2=py38hecd8cb5_0
  - joblib=1.1.1=py38hecd8cb5_0
  - jpeg=9e=hca72f7f_0
  - json5=0.9.6=pyhd3eb1b0_0
  - jsonschema=4.17.3=py38hecd8cb5_0
  - jupyter_client=7.4.9=py38hecd8cb5_0
  - jupyter_core=5.2.0=py38hecd8cb5_0
  - jupyter_server=1.23.4=py38hecd8cb5_0
  - jupyterlab=3.4.4=py38hecd8cb5_0
  - jupyterlab_pygments=0.1.2=py_0
  - jupyterlab_server=2.19.0=py38hecd8cb5_0
  - jupyterlab_widgets=1.0.0=pyhd3eb1b0_1
  - keras=2.9.0=py38hecd8cb5_0
  - keras-preprocessing=1.1.2=pyhd3eb1b0_0
  - kiwisolver=1.4.4=py38hcec6c5f_0
  - krb5=1.19.4=hdba6334_0
  - lcms2=2.12=hf1fd2bf_0
  - lerc=3.0=he9d5cce_0
  - libbrotlicommon=1.0.9=hca72f7f_7
  - libbrotlidec=1.0.9=hca72f7f_7
  - libbrotlienc=1.0.9=hca72f7f_7
  - libcurl=7.84.0=h6dfd666_0
  - libcxx=14.0.6=h9765a3e_0
  - libdb=6.2.32=he9d5cce_1
  - libdeflate=1.8=h9ed2024_5
  - libedit=3.1.20221030=h6c40b1e_0
  - libev=4.33=h9ed2024_1
  - libffi=3.3=hb1e8313_2
  - libgfortran=5.0.0=11_3_0_hecd8cb5_28
  - libgfortran5=11.3.0=h9dfd629_28
  - libiconv=1.16=hca72f7f_2
  - libidn11=1.33=h1de35cc_0
  - libllvm11=11.1.0=h46f1229_6
  - libnghttp2=1.46.0=ha29bfda_0
  - libpng=1.6.37=ha441bb4_0
  - libprotobuf=3.20.1=h8346a28_0
  - libsodium=1.0.18=h1de35cc_0
  - libssh2=1.10.0=h0a4fc7d_0
  - libtiff=4.4.0=h2ef1027_0
  - libwebp=1.2.4=hf6ce154_1
  - libwebp-base=1.2.4=h6c40b1e_1
  - libxml2=2.9.14=hbf8cd5e_0
  - libxslt=1.1.35=h5b33f42_0
  - libzlib=1.2.12=hfd90126_3
  - lightgbm=3.2.1=py38h23ab428_0
  - llvm-openmp=14.0.6=h0dcd299_0
  - llvmlite=0.39.1=py38h8346a28_0
  - lxml=4.9.1=py38h65b224f_0
  - lz4-c=1.9.4=hcec6c5f_0
  - mafft=7.515=h1e3017b_0
  - markdown=3.4.1=py38hecd8cb5_0
  - markupsafe=2.1.1=py38hca72f7f_0
  - matplotlib=3.5.2=py38hecd8cb5_0
  - matplotlib-base=3.5.2=py38hfb0c5b7_0
  - matplotlib-inline=0.1.6=py38hecd8cb5_0
  - minced=0.4.2=hdfd78af_1
  - mistune=0.8.4=py38h1de35cc_1001
  - mkl=2021.4.0=hecd8cb5_637
  - mkl-service=2.4.0=py38h9ed2024_0
  - mkl_fft=1.3.1=py38h4ab4a9b_0
  - mkl_random=1.2.2=py38hb2f4e1b_0
  - multidict=6.0.2=py38hca72f7f_0
  - munkres=1.1.4=py_0
  - muscle=5.1=hb339e23_1
  - nb_conda_kernels=2.3.1=py38hecd8cb5_0
  - nbclassic=0.5.2=py38hecd8cb5_0
  - nbclient=0.5.13=py38hecd8cb5_0
  - nbconvert=6.5.4=py38hecd8cb5_0
  - nbformat=5.7.0=py38hecd8cb5_0
  - ncurses=6.4=hcec6c5f_0
  - nest-asyncio=1.5.6=py38hecd8cb5_0
  - notebook=6.5.2=py38hecd8cb5_0
  - notebook-shim=0.2.2=py38hecd8cb5_0
  - numba=0.56.4=np1.11py3.8hbb42477_g288a38bbd_0
  - numexpr=2.8.4=py38he696674_0
  - numpy=1.21.5=py38h2e5f0a9_3
  - numpy-base=1.21.5=py38h3b1a694_3
  - oauthlib=3.2.1=py38hecd8cb5_0
  - openjdk=11.0.13=h8346a28_0
  - openssl=1.1.1t=hca72f7f_0
  - opt_einsum=3.3.0=pyhd3eb1b0_1
  - packaging=22.0=py38hecd8cb5_0
  - paml=4.9=hb4d813b_6
  - pandas=1.4.4=py38he9d5cce_0
  - pandasql=0.7.3=py38hecd8cb5_1
  - pandocfilters=1.5.0=pyhd3eb1b0_0
  - parallel=20230122=h694c41f_0
  - parso=0.8.3=pyhd3eb1b0_0
  - patsy=0.5.3=py38hecd8cb5_0
  - pcre=8.45=h23ab428_0
  - perl=5.32.1=2_h0d85af4_perl5
  - perl-algorithm-diff=1.201=pl5321hd8ed1ab_0
  - perl-archive-tar=2.40=pl5321hdfd78af_0
  - perl-base=2.23=pl5321hd8ed1ab_0
  - perl-bio-asn1-entrezgene=1.73=pl5321hdfd78af_3
  - perl-bio-coordinate=1.007001=pl5321hdfd78af_3
  - perl-bio-featureio=1.6.905=pl5321hdfd78af_4
  - perl-bio-samtools=1.43=pl5321h1f540d2_3
  - perl-bio-searchio-hmmer=1.7.3=pl5321hdfd78af_0
  - perl-bio-tools-phylo-paml=1.7.3=pl5321hdfd78af_3
  - perl-bio-tools-run-alignment-clustalw=1.7.4=pl5321hdfd78af_3
  - perl-bio-tools-run-alignment-tcoffee=1.7.4=pl5321hdfd78af_5
  - perl-bioperl=1.7.8=hdfd78af_1
  - perl-bioperl-core=1.7.8=pl5321hdfd78af_1
  - perl-bioperl-run=1.007003=pl5321hdfd78af_0
  - perl-business-isbn=3.007=pl5321hd8ed1ab_0
  - perl-business-isbn-data=20210112.006=pl5321hd8ed1ab_0
  - perl-carp=1.50=pl5321hd8ed1ab_0
  - perl-class-data-inheritable=0.09=pl5321h694c41f_0
  - perl-common-sense=3.75=pl5321hd8ed1ab_0
  - perl-compress-raw-bzip2=2.201=pl5321h775f41a_0
  - perl-compress-raw-zlib=2.202=pl5321h775f41a_0
  - perl-constant=1.33=pl5321hd8ed1ab_0
  - perl-data-dumper=2.183=pl5321h775f41a_0
  - perl-db_file=1.858=pl5321hac89ed1_0
  - perl-devel-stacktrace=2.04=pl5321h694c41f_0
  - perl-digest-hmac=1.04=pl5321hdfd78af_0
  - perl-digest-md5=2.58=pl5321hb7f2c08_0
  - perl-encode=3.19=pl5321hb7f2c08_0
  - perl-encode-locale=1.05=pl5321hdfd78af_7
  - perl-exception-class=1.45=pl5321hdfd78af_0
  - perl-exporter=5.74=pl5321hd8ed1ab_0
  - perl-exporter-tiny=1.002002=pl5321hd8ed1ab_0
  - perl-extutils-makemaker=7.66=pl5321hd8ed1ab_0
  - perl-file-listing=6.15=pl5321hdfd78af_0
  - perl-file-path=2.18=pl5321hd8ed1ab_0
  - perl-file-slurp-tiny=0.004=pl5321hdfd78af_2
  - perl-file-sort=1.01=pl5321hdfd78af_3
  - perl-file-spec=3.48_01=pl5321hdfd78af_2
  - perl-file-temp=0.2304=pl5321hd8ed1ab_0
  - perl-getopt-long=2.54=pl5321hdfd78af_0
  - perl-html-parser=3.80=pl5321hb3ea9af_0
  - perl-html-tagset=3.20=pl5321hdfd78af_4
  - perl-http-cookies=6.10=pl5321hdfd78af_0
  - perl-http-daemon=6.14=pl5321hdfd78af_0
  - perl-http-date=6.05=pl5321hdfd78af_0
  - perl-http-message=6.36=pl5321hdfd78af_0
  - perl-http-negotiate=6.01=pl5321hdfd78af_4
  - perl-io-compress=2.201=pl5321h9722bc1_0
  - perl-io-html=1.004=pl5321hdfd78af_0
  - perl-io-socket-ssl=2.075=pl5321hd8ed1ab_0
  - perl-io-string=1.08=pl5321hdfd78af_4
  - perl-io-tty=1.16=pl5321h775f41a_0
  - perl-io-zlib=1.14=pl5321hdfd78af_0
  - perl-ipc-run=20200505.0=pl5321hdfd78af_0
  - perl-json=4.10=pl5321hdfd78af_0
  - perl-json-xs=2.34=pl5321hcd10b59_5
  - perl-libwww-perl=6.67=pl5321hdfd78af_0
  - perl-libxml-perl=0.08=pl5321hdfd78af_3
  - perl-list-moreutils=0.430=pl5321hdfd78af_0
  - perl-list-moreutils-xs=0.430=pl5321ha5712d3_1
  - perl-lwp-mediatypes=6.04=pl5321hdfd78af_1
  - perl-mime-base64=3.16=pl5321h775f41a_0
  - perl-net-http=6.22=pl5321hdfd78af_0
  - perl-net-ssleay=1.92=pl5321hb044416_1
  - perl-ntlm=1.09=pl5321hdfd78af_5
  - perl-parent=0.241=pl5321hd8ed1ab_0
  - perl-pathtools=3.75=pl5321h775f41a_0
  - perl-scalar-list-utils=1.63=pl5321h775f41a_0
  - perl-socket=2.027=pl5321ha5712d3_3
  - perl-storable=3.15=pl5321h775f41a_0
  - perl-sub-uplevel=0.2800=pl5321h775f41a_0
  - perl-test-deep=1.130=pl5321hd8ed1ab_0
  - perl-test-differences=0.69=pl5321h694c41f_0
  - perl-test-exception=0.43=pl5321hd8ed1ab_0
  - perl-test-most=0.38=pl5321hdfd78af_0
  - perl-test-warn=0.37=pl5321hd8ed1ab_0
  - perl-text-diff=1.45=pl5321hd8ed1ab_0
  - perl-time-local=1.30=pl5321hdfd78af_0
  - perl-timedate=2.33=pl5321hdfd78af_2
  - perl-tree-dag_node=1.32=pl5321hdfd78af_0
  - perl-try-tiny=0.31=pl5321h694c41f_0
  - perl-types-serialiser=1.01=pl5321hdfd78af_0
  - perl-uri=5.12=pl5321hdfd78af_0
  - perl-url-encode=0.03=pl5321h527b516_0
  - perl-www-robotrules=6.02=pl5321hdfd78af_4
  - perl-xml-dom=1.46=pl5321hdfd78af_1
  - perl-xml-dom-xpath=0.14=pl5321hdfd78af_2
  - perl-xml-namespacesupport=1.12=pl5321hd8ed1ab_0
  - perl-xml-parser=2.44_01=pl5340h2bc98a8_0
  - perl-xml-regexp=0.04=pl5321hdfd78af_3
  - perl-xml-sax=1.02=pl5321hd8ed1ab_0
  - perl-xml-sax-base=1.09=pl5321hd8ed1ab_0
  - perl-xml-sax-expat=0.51=pl5321hd8ed1ab_0
  - perl-xml-simple=2.25=pl5321hdfd78af_2
  - perl-xml-xpathengine=0.14=pl5321hdfd78af_3
  - pexpect=4.8.0=pyhd3eb1b0_3
  - pickleshare=0.7.5=pyhd3eb1b0_1003
  - pillow=9.2.0=py38hde71d04_1
  - pip=22.3.1=py38hecd8cb5_0
  - pkgutil-resolve-name=1.3.10=py38hecd8cb5_0
  - platformdirs=2.5.2=py38hecd8cb5_0
  - prodigal=2.6.3=ha5712d3_4
  - prokka=1.14.6=pl5321hdfd78af_4
  - prometheus_client=0.14.1=py38hecd8cb5_0
  - prompt-toolkit=3.0.36=py38hecd8cb5_0
  - protobuf=3.20.1=py38he9d5cce_0
  - psutil=5.9.0=py38hca72f7f_0
  - ptyprocess=0.7.0=pyhd3eb1b0_2
  - pure_eval=0.2.2=pyhd3eb1b0_0
  - pyasn1=0.4.8=pyhd3eb1b0_0
  - pyasn1-modules=0.2.8=py_0
  - pycparser=2.21=pyhd3eb1b0_0
  - pygments=2.11.2=pyhd3eb1b0_0
  - pyjwt=2.4.0=py38hecd8cb5_0
  - pyopenssl=22.0.0=pyhd3eb1b0_0
  - pyparsing=3.0.9=py38hecd8cb5_0
  - pyrsistent=0.18.0=py38hca72f7f_0
  - pysocks=1.7.1=py38_1
  - python=3.8.13=hdfd78df_0
  - python-dateutil=2.8.2=pyhd3eb1b0_0
  - python-fastjsonschema=2.16.2=py38hecd8cb5_0
  - python-flatbuffers=1.12=pyhd3eb1b0_0
  - python_abi=3.8=2_cp38
  - pytz=2022.7=py38hecd8cb5_0
  - pyyaml=6.0=py38h6c40b1e_1
  - pyzmq=23.2.0=py38he9d5cce_0
  - re2=2022.04.01=he9d5cce_0
  - readline=8.2=hca72f7f_0
  - requests=2.28.1=py38hecd8cb5_0
  - requests-oauthlib=1.3.0=py_0
  - rsa=4.7.2=pyhd3eb1b0_1
  - scipy=1.7.3=py38h214d14d_2
  - seaborn=0.12.2=py38hecd8cb5_0
  - send2trash=1.8.0=pyhd3eb1b0_1
  - setuptools=65.6.3=py38hecd8cb5_0
  - shap=0.41.0=py38h07fba90_0
  - six=1.16.0=pyhd3eb1b0_1
  - slicer=0.0.7=pyhd3eb1b0_0
  - snappy=1.1.9=he9d5cce_0
  - sniffio=1.2.0=py38hecd8cb5_1
  - soupsieve=2.3.2.post1=py38hecd8cb5_0
  - sqlalchemy=1.4.39=py38hca72f7f_0
  - sqlite=3.39.3=h707629a_0
  - stack_data=0.2.0=pyhd3eb1b0_0
  - statsmodels=0.13.2=py38hca72f7f_0
  - t-coffee=12.00.7fb08c2=h26a2512_0
  - tbl2asn-forever=25.7.2f=ha5712d3_2
  - tensorboard=2.9.0=py38hecd8cb5_0
  - tensorboard-data-server=0.6.1=py38h7242b5c_0
  - tensorboard-plugin-wit=1.6.0=py_0
  - tensorflow=2.9.1=eigen_py38hf4d42e0_1
  - tensorflow-base=2.9.1=eigen_py38h36d2db2_1
  - tensorflow-estimator=2.9.0=py38hecd8cb5_0
  - termcolor=2.1.0=py38hecd8cb5_0
  - terminado=0.17.1=py38hecd8cb5_0
  - threadpoolctl=2.2.0=pyh0d69192_0
  - tinycss2=1.2.1=py38hecd8cb5_0
  - tk=8.6.12=h5d9f67b_0
  - tornado=6.2=py38hca72f7f_0
  - tqdm=4.64.1=py38hecd8cb5_0
  - traitlets=5.7.1=py38hecd8cb5_0
  - typing-extensions=4.4.0=py38hecd8cb5_0
  - typing_extensions=4.4.0=py38hecd8cb5_0
  - urllib3=1.26.14=py38hecd8cb5_0
  - viennarna=2.5.1=py38pl5321h6a1f747_0
  - wcwidth=0.2.5=pyhd3eb1b0_0
  - webencodings=0.5.1=py38_1
  - websocket-client=0.58.0=py38hecd8cb5_4
  - werkzeug=2.2.2=py38hecd8cb5_0
  - wheel=0.35.1=pyhd3eb1b0_0
  - widgetsnbextension=3.5.2=py38hecd8cb5_0
  - wrapt=1.14.1=py38hca72f7f_0
  - xz=5.2.10=h6c40b1e_1
  - yaml=0.2.5=haf1e3a3_0
  - yarl=1.8.1=py38hca72f7f_0
  - zeromq=4.3.4=h23ab428_0
  - zipp=3.11.0=py38hecd8cb5_0
  - zlib=1.2.12=h4dc903c_3
  - zstd=1.5.2=hcb37349_0
  - pip:
      - appdirs==1.4.4
      - et-xmlfile==1.1.0
      - flowcal==1.3.0
      - frozendict==2.3.7
      - html5lib==1.1
      - imageio==2.22.1
      - markdown-it-py==2.1.0
      - matplotx==0.3.10
      - mdurl==0.1.2
      - multitasking==0.0.11
      - networkx==2.8.7
      - openpyxl==3.0.10
      - perfplot==0.10.2
      - pywavelets==1.4.1
      - rich==13.2.0
      - scikit-image==0.19.3
      - scikit-learn==1.1.2
      - tensorflow-addons==0.18.0
      - tifffile==2022.10.10
      - typeguard==2.13.3
      - xlrd==1.2.0
      - yfinance==0.2.18
prefix: /Users/EthanJ/opt/anaconda3/envs/plotting
```
