## Supplementary figures and images for "LanTERN: a fluorescent sensor that specifically binds lanthanides"

### supplemental_figure5.png

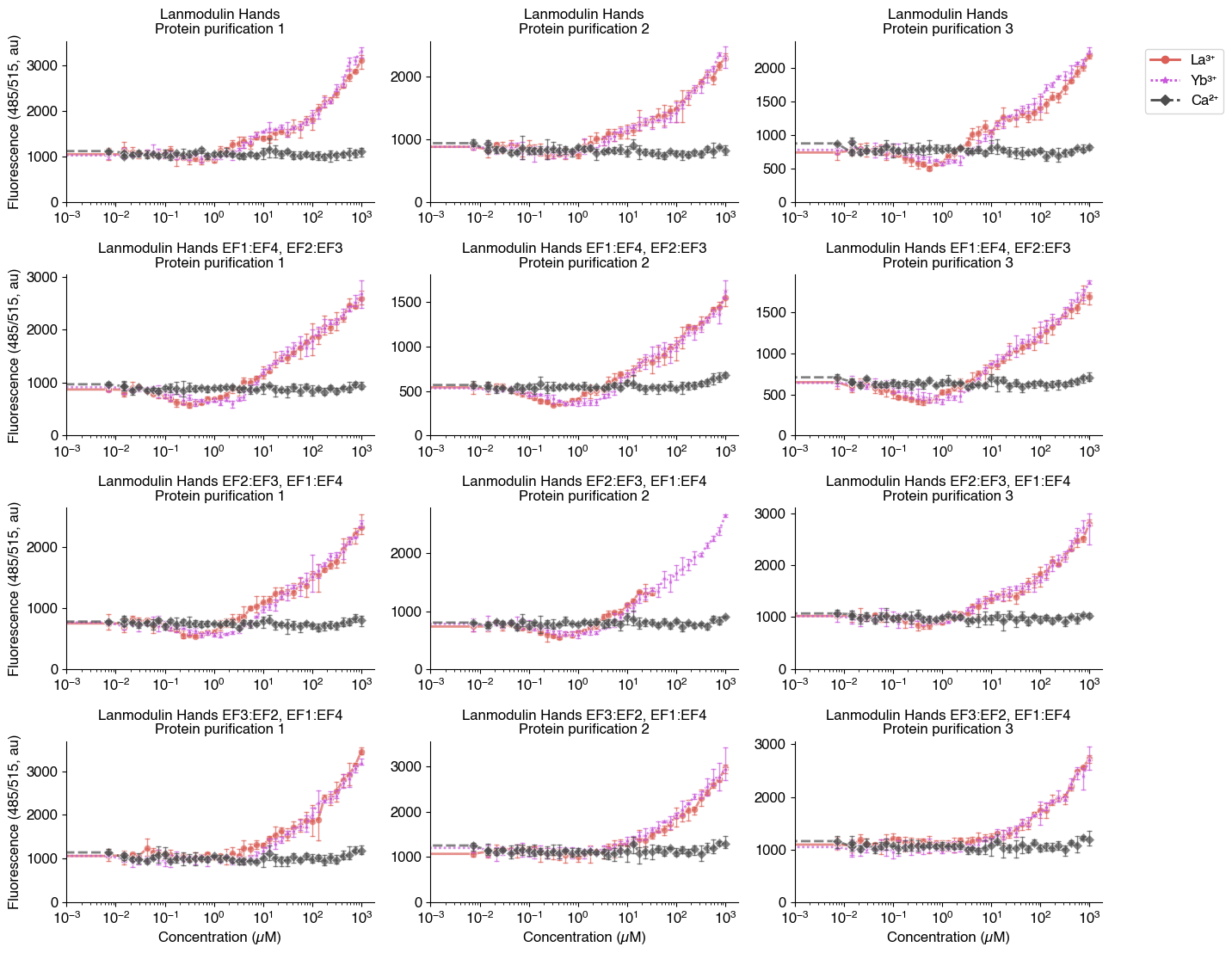
